# Mitochondrion-IMC contact sites are critical for cofactor biosynthesis and egress signaling in *Toxoplasma gondii*

**DOI:** 10.64898/2026.04.08.717193

**Authors:** Rodolpho Ornitz Oliveira Souza, Katherine Thibodeau, Kylie Jacobs, Chunlin Yang, Marta T. Gomes, Gustavo Arrizabalaga

## Abstract

*Toxoplasma gondii* is a single-celled parasite belonging to the Apicomplexa phylum. *Toxoplasma*’s single mitochondrion is highly dynamic, changing its morphology as the parasite undergoes egress and invasion. Recently, we have demonstrated that mitochondrial morphology is driven by a protein named Lasso Maintenance Factor 1 (LMF1). This protein interacts with IMC10, a protein present at the parasite’s inner membrane complex (IMC), mediating a unique membrane contact site between the IMC and mitochondrion. Interestingly, parasites lacking either LMF1 or IMC10 have abnormal mitochondrial morphology, cell division defects, and delayed propagation in tissue culture. Although both components of the tether were identified, the functions of this contact site remain unknown. In this work, we show that Δ*lmf1* parasites exhibit upregulation of egress signaling and downregulation in folate metabolism and pantothenate biosynthesis. Δ*lmf1* parasites exhibit increased intracellular calcium levels, leading to greater sensitivity to ionophore-induced egress and microneme secretion. We have confirmed that parasites have decreased levels of tetrahydrofolate and coenzyme A, showing a limitation in cofactor production. Interestingly, the Δ*lmf1* parasites prefer glutamine instead of glucose as a catabolic substrate. Accordingly, we demonstrate for the first time that proper mitochondrial positioning is crucial for folate and Coenzyme A metabolism as well as egress signaling.

**IMPORTANCE:** *Toxoplasma gondii* is the causative agent of Toxoplasmosis, a disease that affects a third of the world’s population. This parasite has a single, highly dynamic mitochondrion. The parasite’s mitochondrion changes shape depending on environmental conditions (inside or outside the host cell) or on stressors, such as drugs. Our laboratory characterized the proteins involved in regulating mitochondrial dynamics in the parasite, but the functional importance of these mitochondrial changes has not yet been described. Here, we show that the shape of Toxoplasma’s mitochondrion is important for the synthesis of key cofactors, such as folates and coenzyme A. We show that mitochondrial shape in this parasite is important for signaling the parasite’s exit from the host cell, a critical process in its life cycle. These findings review a previously unknown function of a parasite-specific organelle contact site, providing new insights into the importance of mitochondria for these parasites.

## INTRODUCTION

*Toxoplasma gondii* is an obligatory intracellular parasite that belongs to the Apicomplexa phylum [1]. It can infect a wide range of warm-blooded animals, including humans [2]. Although most human infections are asymptomatic, toxoplasmosis can result in severe disease and death in immunosuppressed individuals [3] and those infected congenitally [4]. Drug treatments for this pathogen are limited to a few options that are often toxic and are ineffective against the chronic form of the parasite. Therefore, expanding the repertoire of therapeutic targets in *Toxoplasma* is a priority.

A validated target for drug development is the parasite’s single mitochondrion, which is essential for parasite survival [5]. While much is known about the biochemistry of the *Toxoplasma* mitochondrion, relatively little is known about the mechanisms and factors that regulate its structure and inheritance. *Toxoplasma*’s tubular mitochondrion is highly dynamic, exhibiting diverse morphologies during the parasite’s lytic cycle [6, 7] and in response to stressors [8]. When inside its host cell, *Toxoplasma* presents a lasso-shaped mitochondrion that spans the parasite’s periphery. Upon egress, the mitochondrion rapidly detaches from the pellicle, transitioning first to a sperm-like morphology and finally collapsing towards the apical end of the parasite. Soon after the parasite invades another host cell, the lasso shape is reconstituted. When in the lasso morphology, the mitochondrion abuts the parasite’s inner membrane complex (IMC) [6, 9], establishing what is reminiscent of membrane contact sites. The functional relevance of the highly coordinated positioning of the mitochondrion is unclear, but the identification of proteins responsible for tethering the organelle to the IMC suggests that it is important for parasite fitness.

We previously described a mitochondrial outer membrane-localized protein, LMF1, critical for mitochondrial positioning at the parasite’s periphery [7]. Loss of LMF1 causes the mitochondrion of intracellular parasites to collapse, akin to what is only normally seen in extracellular parasites. Using protein-protein interaction techniques and ultrastructure expansion microscopy (U-ExM), we identified IMC10, a pellicle-localized protein, as the IMC side of the tether, cooperating with LMF1 in mitochondrial positioning and distribution [10]. Disruption of either LMF1 or IMC10 leads to defects in mitochondrial inheritance, suggesting a role of this tether in mitochondrial division [7, 10]. In addition, parasites lacking LMF1 exhibit a propagation phenotype, indicating that proper mitochondrial positioning is important for propagation in tissue culture [7]. As we gain a better understanding of the protein composition of this IMC-mitochondrion tether, we can explore the functionality of this contact site. Our work brings the first insights into the physiological importance of mitochondrial dynamics in *Toxoplasma*. Here, we demonstrate that the loss of LMF1 results in decreased Dihydrofolate Reductase-Thymidylate synthetase (DHFR-TS) activity and impaired Coenzyme A biosynthesis. These mutant parasites have higher mitochondrial metabolism and a shift towards glutamine as their preferred catabolic source. Intriguingly, the loss of contact sites results in higher intracellular calcium levels, promoting more efficient egress and differences in microneme positioning and secretion. Thus, mitochondrial positioning and morphology appear to be critical for parasite metabolism and signaling.

## RESULTS

### Loss of LMF1 affects the transcription of metabolic and egress signaling-related genes

To determine the functional relevance of the molecular tether between the mitochondrion and the IMC, we investigated the effect of losing LMF1, and consequently the mitochondrion-IMC tethering, on the parasite’s transcriptome. We evaluated the levels of transcripts by RNA sequencing using mRNA extracted from parental (Δ*ku80*) and the previously established Δ*lmf1* strain [7] (Supplemental dataset 1). Genes of significance were identified from triplicate assays based on an adjusted p-value < 0.05 and a log2 score >1 or <-1 (Table 1). To highlight dysregulated genes, the results were displayed in a volcano plot (Fig. 1). Comparing the transcriptome of the Δ*lmf1* parasites with that of the parental strain, we observed that 109 genes were differentially expressed. Of these, 35 genes showed decreased transcript levels, and 74 increased transcript levels. As expected, *LMF1* exhibited a negative Log2 fold change, confirming its disruption. Of note, the expression of IMC10, the interacting protein that mediates tethering, was not significantly affected by the absence of LMF1 (Supplemental dataset 1).

**Figure 1.**
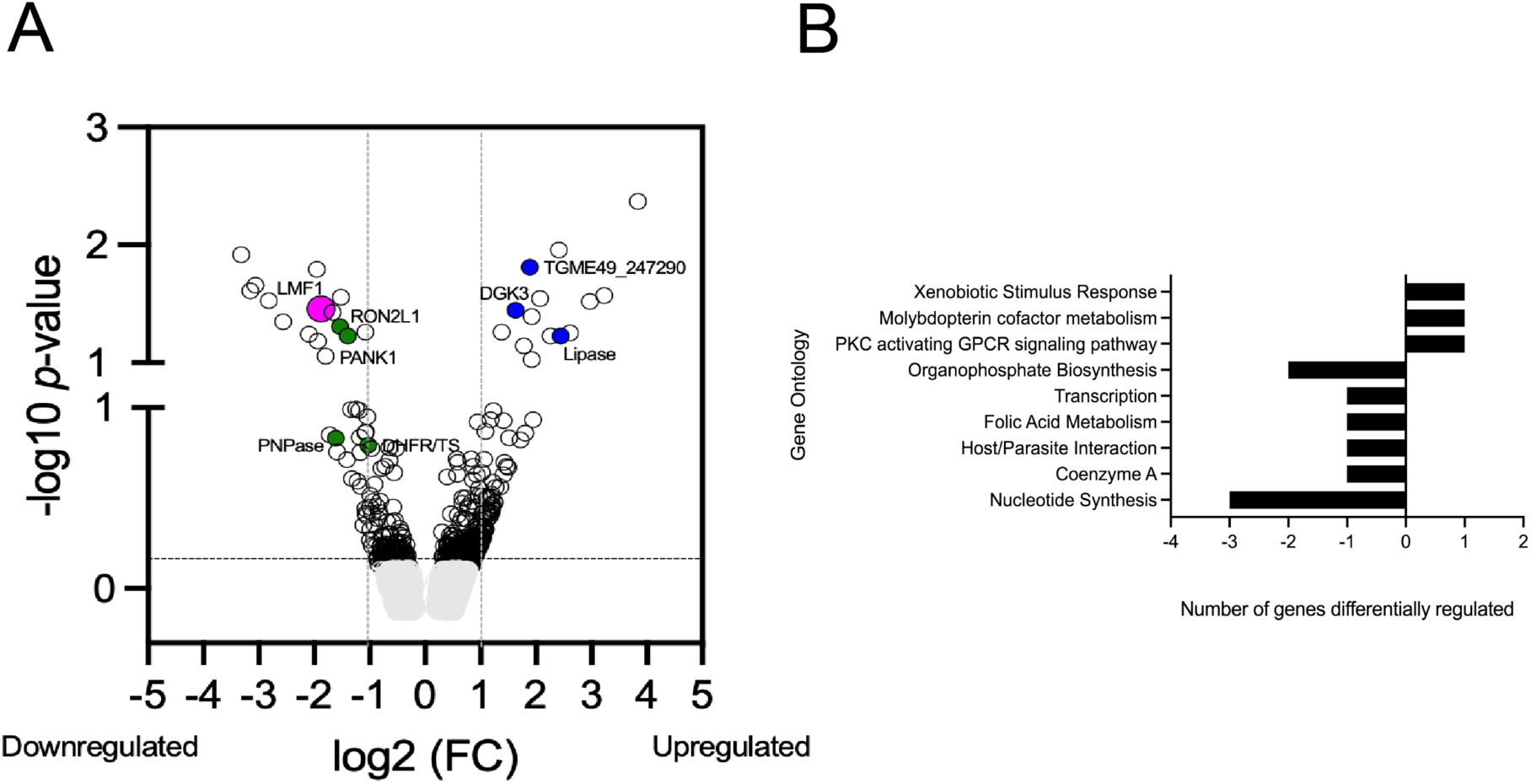
RNA sequencing shows differential gene expression in the LMF1-KO parasites. A) Volcano plot showing all differentially expressed genes with a log2 Fold Change >0.5 and p < 0.05 in Δ*lmf1* parasites. Genes of interest are highlighted with colored circles: pink = LMF1; green = downregulated genes of interest; blue = upregulated genes of interest. The experiment was performed in triplicate. B) Bar graph showing the Gene Ontology (GO) for the main pathways up- and downregulated in the Δ*lmf1*.

**Table 1.**
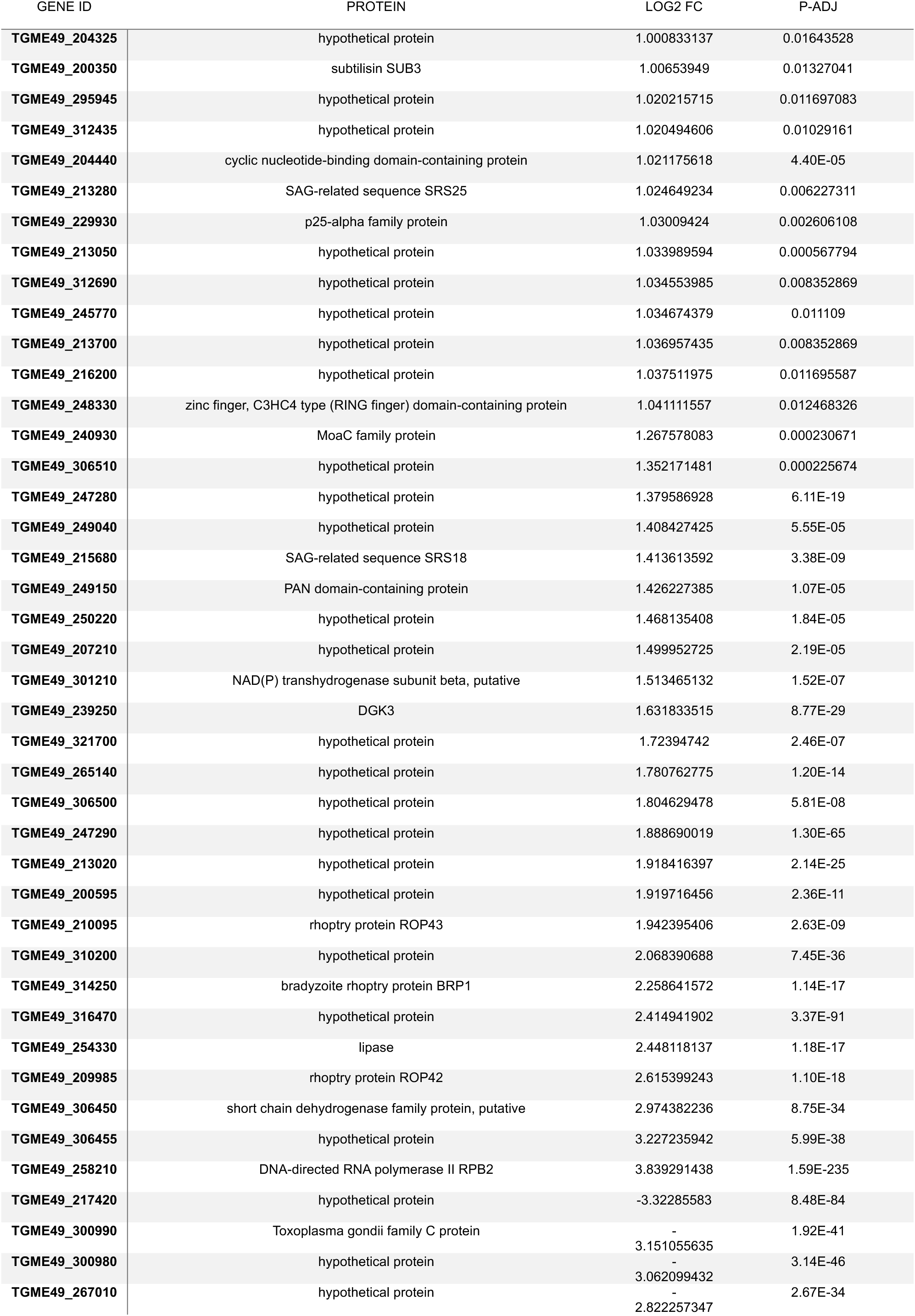
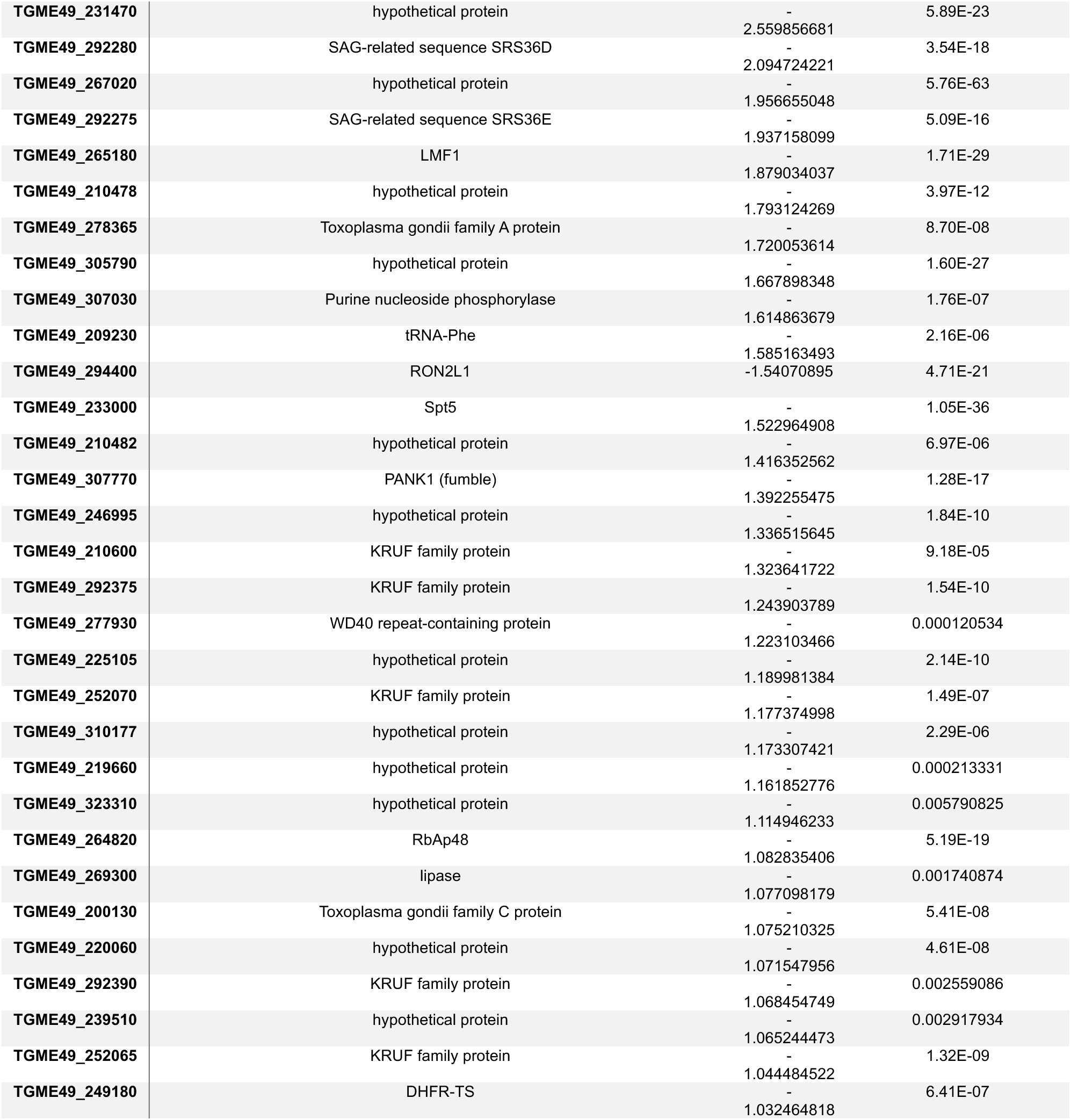
Summary of differentially regulated genes in the Δ*lmf1* strain. All differentially expressed genes with a log2 Fold Change >0.5 and p < 0.05 in Δ*lmf1* parasites. The full dataset is available at Supplemental dataset 1.

The 109 differentially expressed genes were analyzed using Gene Ontology enrichment [11] to identify their corresponding pathways (Fig. 1B). Our analyses showed that two metabolic genes were downregulated: pantothenate kinase 1 (Pank1; TGME49_307770) and dihydrofolate reductase-thymidylate synthase (DHFR-TS; TGME49_249180). Pank1 is the first cytosolic step of Coenzyme A (CoA) biosynthesis in *Toxoplasma*. Alveolates possess a unique polypeptide for the bifunctional DHFR-TS, which connects folate and nucleotide metabolism [12]. DHFR-TS is the main target of pyrimethamine, a drug used to treat toxoplasmosis and malaria [13]. The precise subcellular location of DHFR-TS in *Toxoplasma* is unclear, as previous spatial proteomics (HyperLOPIT) data have detected it in both the nucleus and the cytosol [14, 15]. Other genes that appeared to be downregulated include those involved in nucleotide metabolism, such as purine nucleoside phosphorylase (PNPase, TGME49_307030) [16] and RON2L1 (TGME49_294400), a protein linked to invasion processes.

Among the upregulated genes (Table 1, Supplemental dataset 1) are those related to xenobiotic response stimuli (Tartrolon E responsive gene, TGME49_272370) [17], molybdopterin cofactor metabolic process (MoaC family protein, TGME49_240930), and genes that are related to lipid metabolism and second messenger, such as a putative lipase (TGME49_254330), a putative acylglycerol lipase (TGME49_247290), and diacylglycerol kinase 3 (DGK3, TGME49_239250). *Toxoplasma* possesses three DGKs (DGK1, 2, and 3), and DGK3 is a non-essential protein for parasite survival [18]. These proteins are involved in egress signaling and are critical for triggering microneme secretion and parasite motility [18, 19].

To confirm that the differential expression of genes in the knockout strain was specifically related to the loss of LMF1, we assessed the global transcriptomics of a knockout strain that has been complemented with an exogenous copy of LMF1 (Δ*lmf1*+*LMF1*-HA) [7] (Supplemental fig. S1, Supplemental dataset 2). Comparing mRNA levels in the complemented strain to those in the Δ*lmf1* and parental strains reveals that genes dysregulated in the knockout strain exhibit levels like those of the parental strain. Of note, *LMF1* mRNA levels in the complemented strain are lower than those of the endogenous copy in the parental strain (Supplemental dataset 3). The reduced expression of *LMF1* in the complemented strain is consistent with the observation that the mitochondrial morphology phenotype is only partially rescued in this strain [7]. Similarly, we observe that expression of some genes, such as Pank1, is only partially recovered, confirming that this gene’s expression may be related to *LMF1* levels and/or the mitochondrial morphology (Supplemental fig. S1, Supplemental dataset 3).

### Δ*lmf1* parasites exhibit enhanced sensitivity to induced egress

The RNA-seq data showed that DGK3 transcript levels were higher in the Δ*lmf1* strain. DGK proteins mediate the synthesis of diacylglycerol (DAG) to phosphatidic acid (PA), and vice versa (Fig. 2A). PA plays an important role in egress, and *Toxoplasma* can both synthesize it or uptake it from the host [19]. Thus, it is plausible that an inability to take up PA, due to the altered mitochondrial positioning, increases reliance on DGK synthesis, leading to its upregulation. Accordingly, we assessed PA uptake by incubating freshly syringe-released parasites with a fluorescent probe, 16:0-06:0 NBD PA (ePA-488) [20] for 10 minutes. Parasites were then stained for the surface antigen SAG1 using a standard immunofluorescence assay. Parental parasites effectively took up probe, which appears to be dispersed throughout the parasite with no specific compartmentalization (Fig. 2B). The same pattern of staining was observed for the Δ*lmf1* parasites (Fig. 2B). We quantified the total fluorescence of PA staining and divided it by the fluorescence intensity of Sag1 (ratio ePA/SAG1) (Fig. 2B). No statistical difference was observed when comparing both strains, showing that the mutant parasites can take up PA from the external media with the same efficiency as the parental ones. In conclusion, there appears to be no correlation between DGK3 levels and PA uptake in the absence of LMF1.

**Figure 2.**
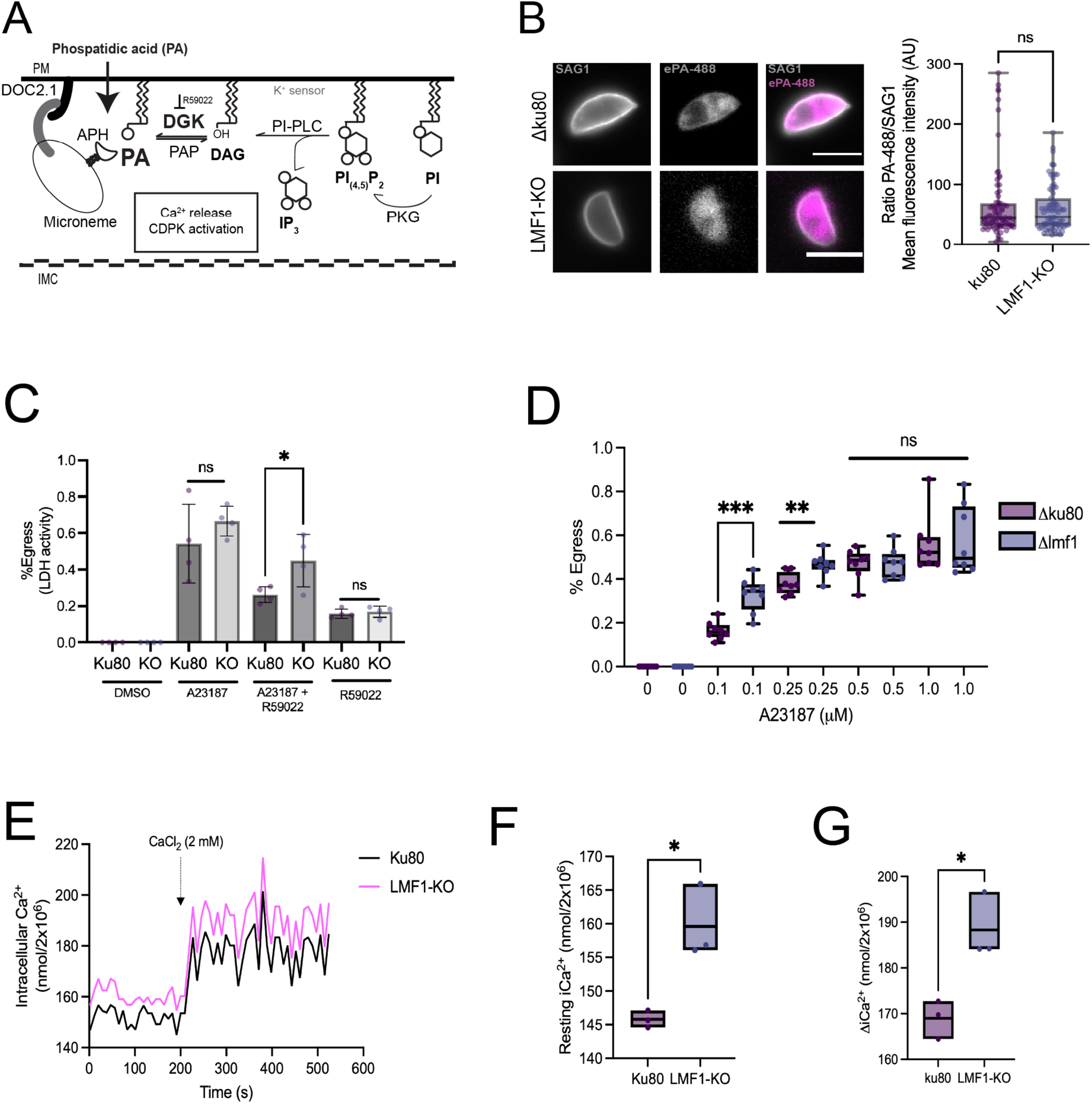
Loss of LMF1 facilitates Ca^2+^-induced egress. A) Schematic representation of Ca^2+^ and lipid second messenger signaling in *Toxoplasma*. B) External phosphatidic acid (ePA) uptake assay. Syringe-released parasites were incubated with PA-488 for 1 min, then fixed and stained with anti-SAG1. Individual parasites were imaged and then used to measure the mean fluorescence intensity ratio (ePA/SAG1). C) Measurement of egress induced by 2 µM A2318. Egress was measured by the levels of lactate dehydrogenase released from the host cells upon parasite bursting. D) Parasite egress using different concentrations of A23187. E) Intracellular parasites from the parental and knockout strains were loaded with Fura-2 AM to monitor calcium levels. The arrow represents the time at which 2 mM CaCl_2_ was spiked into the medium. F) Resting levels of intracellular CaCl_2_ from both parental and KO cell lines. All data graphs are the average of three biological triplicates (n=3). Data are means ± s.d.. Statistical analysis is a Two-tailed unpaired t-test, ** P<0.01; * P<0.05, ns = not significant.

Because DGK mRNA levels are increased in the absence of LMF1, we tested whether the knockout strain exhibited altered sensitivity to the DGK inhibitor R59022, which inhibits parasite Ca^2+^-stimulated egress and microneme secretion [18]. For this purpose, we induced egress in the knockout and parental strains with the calcium ionophore A23187 in the presence or absence of R59022. Parasite egress was evaluated by measuring lactate dehydrogenase activity in the media, which is released from the host cell upon parasite exit. The parental and knockout parasites undergo egress at similar levels when incubated with 1 µM A23187 (Fig. 2C). As expected, the addition of R59022 inhibits A23187-induced egress in the parental strain (Fig. 2C). By contrast, the Δ*lmf1* parasites are less sensitive to DGK inhibition (Fig. 2C), which is likely due to the higher levels of DGK.

In addition to the observed reduced susceptibility to R59022 inhibition, an increase in DGK protein levels might render the parasite more susceptible to egress induction. Accordingly, we tested egress levels of the parental and Δ*lmf1* and parental strains at suboptimal concentrations of A23187 (Fig. 2D). Both parasite strains exhibited a concentration-dependent sensitivity to A23187-induced egress (Fig. 2D). Interestingly, the knockout parasites exhibited statistically significantly higher rates of egress than the parental strain at the lower concentrations of the ionophore (0.1 and 0.25 µM), supporting the hypothesis that the Δ*lmf1* strain is more sensitive to induced egress (Fig. 2D). To investigate whether this heightened sensitivity to A23187-induced egress is accompanied with a calcium dysregulation within the parasites, we measured Ca^2+^ levels in the parental and knockout strains. Parasites from both strains were loaded with Fura2-AM, and the levels of intracellular Ca^2+^ (resting) and buffered calcium were measured for 8 minutes (Fig. 2E). In the knockout parasites, the resting Ca^2+^ levels were higher than those in the parental cell lines (Figs. 2E and F) (156.8±5.5 nmols in the knockouts vs 145±1.3 nmols in the parental cell lines). To test whether Ca^2+^ uptake was also affected, we added 2 mM CaCl_2_ to each cell line and observed a significant increase in Ca^2+^ uptake efficiency in the mutant strain (Fig. 2E and G). Taken together, our results demonstrate that the absence of LMF1 and the subsequent loss of proper mitochondrial positioning impact the parasite’s intracellular Ca^2+^ levels and egress.

### Microneme docking and secretion are altered in the LMF1-KO parasites

DGK regulates microneme secretion, a Ca^2+^-dependent process critical for parasite motility and egress [21, 22]. Given that the Δ*lmf1* strain overexpresses DGK and exhibits dysregulated calcium and egress levels, we investigated microneme secretion in the mutant. Normally, micronemes are distributed throughout the cytosol and migrate towards the apical end, from which they secrete their contents [23]. Accordingly, we first analyzed microneme positioning by immunofluorescence using antibodies against the microneme protein MIC5. As expected, we observe micronemes spread throughout the parasite in the parental strain, with some concentration towards the apical end (Fig. 3A). In Δ*lmf1* parasites, the microneme distribution appears different, with a higher concentration of the organelle towards the apical end of the parasite relative to the parental strain (Fig. 3A). This could be confirmed by quantifying the fluorescence intensity at the parasite apical end (Fig. 3A). This higher signal intensity in the knockout strain is not due to an increase in MIC5 expression, as we observed similar protein levels between strains by western blot (Supplemental fig. S2). Thus, it appears that in the absence of LMF1, there is higher accumulation of micronemes towards the apical end of the parasite.

**Figure 3.**
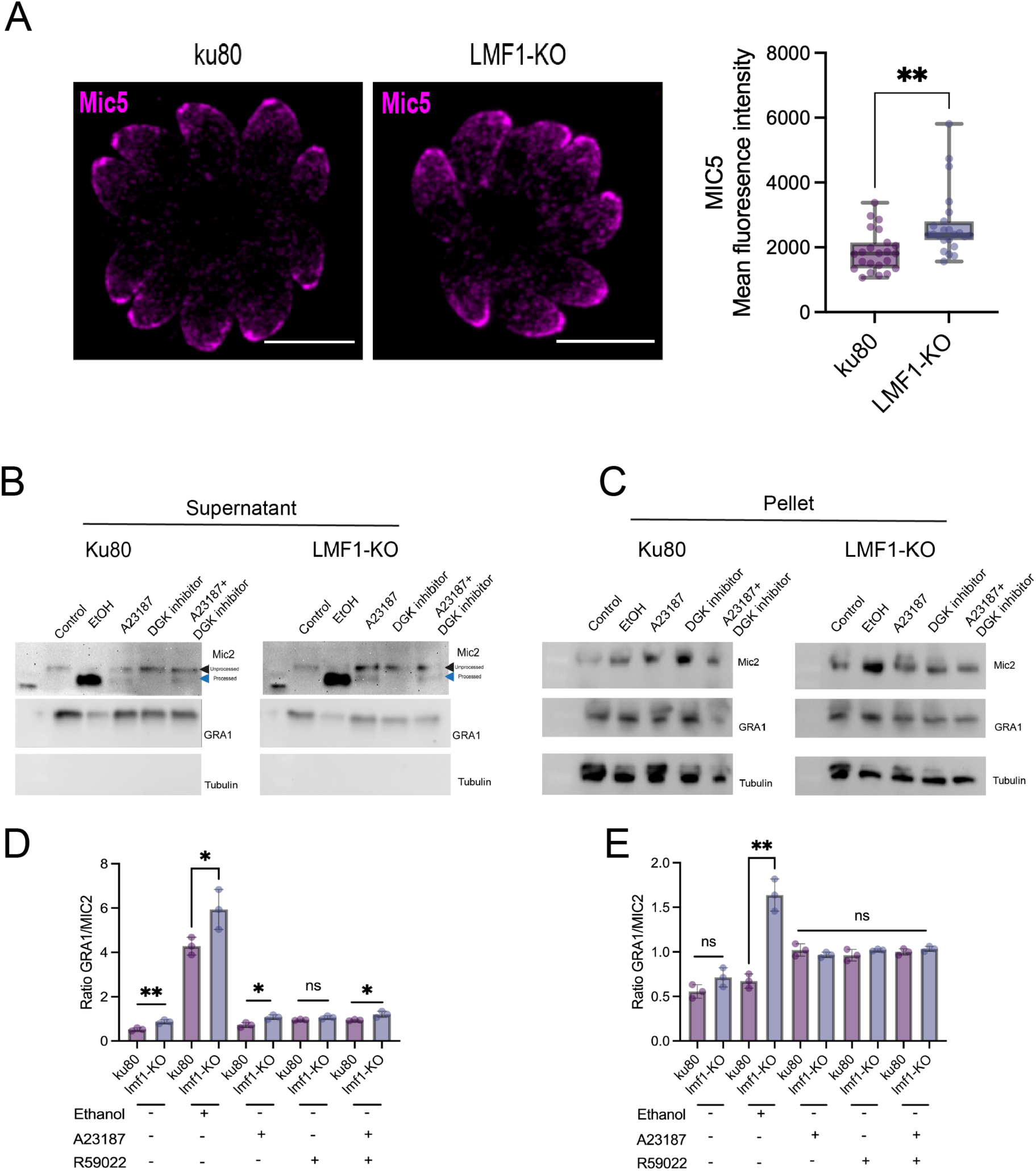
Loss of LMF1 affects microneme docking and secretion. A) Immunofluorescence of parasites stained for MIC5 to visualize micronemes. Intracellular parasites were fixed and stained with an anti-MIC5 antibody. The graph represents fluorescence intensity at the apical end for 35 parasites. The experiment was performed in triplicates (n=3). Data are means ± s.d.. Statistical analysis is a Two-tailed unpaired t-test, ** P<0.01. B) Western blots of supernatant (secreted antigens) and C) pellet (whole parasites) from parental (Ku80) and mutant (Δ*lmf1*) strains. Microneme secretion was induced with 2% EtOH or 2 µM A23187 with or without 30 µM of DGK inhibitor. Parasites were also treated with 30 µM of DGK inhibitor alone. Blots were probed for the microneme protein MIC2 [22], the dense granule protein GRA1 [64], and tubulin [58] (to confirm an equal number of parasites across samples). Arrows in the figure correspond to the unprocessed and processed forms of MIC2. D and E) Densitometry of the levels of proteins from supernatant (D) and pellet (E). Densitometry was calculated by dividing the processed MIC2 intensity ratio by GRA1. Densitometry analyses were performed using FIJI. The experiment was performed in triplicate (n=3). Data are means ± s.d. Statistical analysis is a Two-tailed unpaired t-test, ** P<0.01; * P<0.05, ns = not significant.

To determine whether the concentration of micronemes at the apical end correlates with higher microneme secretion in the knockout, we compared the levels of secreted MIC2 between the parental and Δ*lmf1* strains (Fig. 3B). For this purpose, we syringe-released intracellular parasites and incubated them in plain media (control), 2% EtOH (v/v), 1 µM A23187, or 1 µM A23187 combined with 30 µM of R50922 (Fig. 3C and 3D). Both EtOH and A23187 are potent inducers of microneme secretion [22]. As an additional control, we performed the assay with 30 µM R50922 alone. Levels of the microneme protein MIC2 secreted into the media were analyzed by Western blot. We also determined protein levels from the pelleted parasites after the incubation. It is important to note that microneme proteins such as MIC2 are proteolytically processed at their C-terminus during trafficking and secretion [24]. In non-induced parasites, two bands can be observed, corresponding to processed (lower band) and unprocessed (upper band). Upon induction, a single band corresponding to processed MIC2 is observed. As expected, non-induced parasites of the parental strain secreted negligible amounts of MIC2 into the supernatant. By contrast, the Δ*lmf1* parasites secreted significant levels of MIC2 into the supernatant without induction (Fig. 3B, quantified in 3D). Similarly, in the presence of 2% EtOH or A23187, the Δ*lmf1* parasites secreted more protein into the supernatant relative to the parental strain (Fig. 3B-D). To test whether the observed effects are specific to the microneme, we monitored dense granule secretion, which is not Ca^2+^-dependent. As shown in Fig. 3B and 3C, we observe no differences in GRA1 secretion among the strains tested. Since the knockouts showed reduced sensitivity to the DGK inhibitor R50922, we then tested whether a similar effect was observed in the context of microneme secretion. We first measured microneme secretion in both cell lines in the presence of the inhibitor alone and quantified MIC2 levels in the supernatant. There were no significant differences between the strains in this condition (Fig. 3B-D). When A23187 was combined with R50922, the parental strain parasites still secreted MIC2, but at lower levels than the Δ*lmf1* strain, consistent with observations in the egress-based assays. Taken together, our results indicate that Δ*lmf1* parasites are more efficient at stimulated microneme secretion, likely due to increased DGK3 expression.

### Loss of LMF1 affects DHFR-TS activity

To determine whether the downregulation of the *DHFR-TS* transcript correlates with reduced function, we measured DHFR-TS enzymatic activity in crude extracts of intracellular parasites from both the parental and knockout strains. Briefly, parasites were syringe-released from host cells and purified by filtration to remove host material after syringe lysis. DHFR-TS activity was measured by observing the oxidation of NADPH to NADP^+^ at 340 nm. Subsequently, the absorbance was converted to specific activity (units of enzyme/mg of protein). The Δ*lmf1* parasites exhibit a reduction of 75% in DHFR-TS activity as compared to the parental strain, indicating that the downregulation of *DHFR-TS* transcript in these parasites correlates with a reduction in protein activity (Fig. 4B). We also measured the DHFR-TS activity in the complemented parasite line, which exhibited a level of enzyme activity comparable to that of the parental cell line (Fig. 4B).

**Figure 4.**
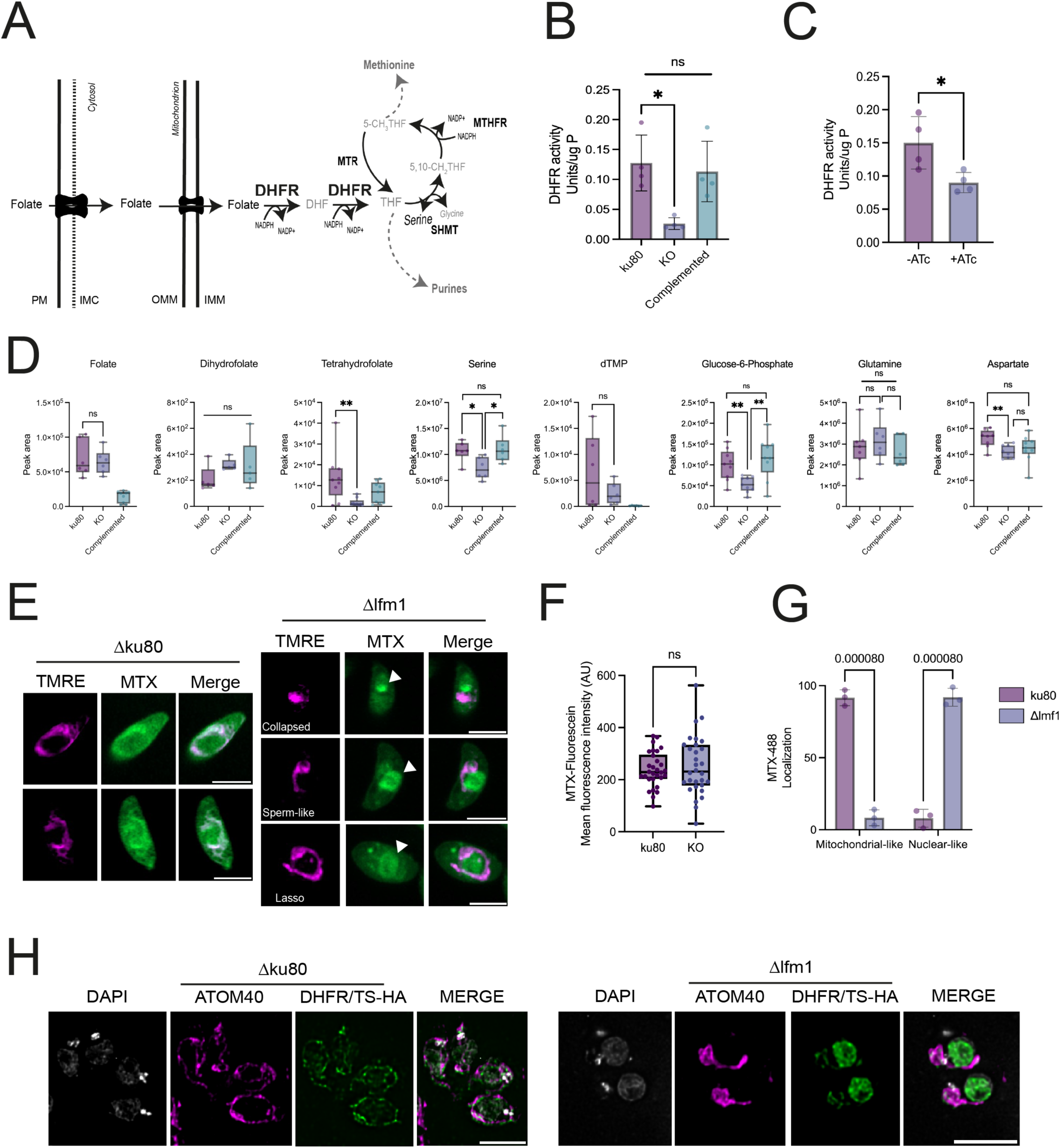
DHFR-TS-TS activity is affected by LMF1 loss. A) Schematic representation of folate metabolism in *Toxoplasma*. PM = plasma membrane. IMC = inner membrane complex. OMM = outer mitochondrial membrane. IMM = inner mitochondrial membrane. DHF = dihydrofolate. DHFR-TS = dihydrofolate reductase. THF = tetrahydrofolate. B) Total activity of DHFR-TS in crude extracts of intracellular tachyzoites. Enzymatic activity was performed in n=4. Data are means ± s.d. Statistical analysis is a Two-tailed unpaired t-test, ** P<0.01. C). Total activity of DHFR-TS in crude extracts of intracellular parasites TATi-IMC10. Parasites were maintained in the presence or absence (control) of anhydrotetracycline (ATc) for 72 hours. Enzymatic activity was performed in n=4. Data are means ± s.d. Statistical analysis is a Two-tailed unpaired t-test, ** P<0.01. D) Untargeted metabolomics results for key metabolites in parasites’ folate pathway. Metabolite quantification was performed in three technical replicates and three biological replicates (n=9). Data are means ± s.d.. Statistical analysis is a Two-tailed unpaired t-test, ** P<0.01; * P<0.05. E) Representative images of parental and knockout parasites after methotrexate-fluorescein (MTX-488) incorporation. Parasites were stained with TMRE to track mitochondria. White arrows indicate a region of the cell that resembles the nucleus. F) Total fluorescence of MTX-488 parasites. Fluorescence intensity was measured in 30 individual parasites (n = 3). G) Quantification of the differential accumulation of MTX-488 in the mitochondrion and the nucleus in parental (ku80) and knockout parasites. A total of 30 individual parasites were compared in each replicate (n=3). Data are means ± s.d. Statistical analysis is a Two-tailed unpaired t-test; ns, not significant (P > 0.05). H) IFA of intracellular parental (Δku80) and knockout (Δlmf1) parasites expressing DHFR-TS with an HA epitope tag. Parasites were stained with DAPI (nuclear marker) and TOM40 (mitochondrial marker).

The loss of LMF1 results in a loss of the tether between the mitochondrion and the inner membrane complex. To determine whether the reduction in DHFR-TS activity is related to the loss of this tethering, we measured DHFR-TS activity in the absence of IMC10, which interacts with LMF1 and is also required to establish the mitochondrion-IMC contact sites [10]. The conditional IMC10 knockdown cell line was grown in the presence of anhydrotetracycline (ATc) for 72 hours, sufficient time to achieve more than 90% reduction in protein expression [10]. We tested DHFR-TS activity in crude extracts from parasites kept in both the presence (+ATc) and absence (DMSO) of ATc. The IMC10 knockdown demonstrated a 34% decrease in DHFR-TS activity compared to the control conditions (DMSO) (Fig. 4C). Together, these results suggest that the tethering of the mitochondrion to the IMC is important for DHFR-TS activity.

### Loss of LMF1 affects folate metabolism and its related metabolites

DHFR-TS metabolizes folate into dihydrofolate and tetrahydrofolate in a NADPH-dependent manner, which in turn feeds into purine, pyrimidine, and methionine biosynthesis (Fig. 4A). In *Toxoplasma*, DHFR-TS is also responsible for the synthesis of dTMP, a pyrimidine nucleotide. To assess whether the lack of LMF1 and concurrent reduction in DHFR-TS function affect downstream metabolic steps, we performed label-free targeted metabolomics to detect changes in the folate cycle and related pathways. We were able to detect folic acid, dihydrofolate (DHFR-TS substrate), tetrahydrofolate (DHFR-TS product), serine (substrate for serine hydroxymethyltransferase, SHMT), deoxythymidine monophosphate (dTMP), glutamine and aspartate (substrates for the de novo pyrimidine biosynthesis), and glucose-6-phosphate (which will then be converted into ribulose-5-phosphate, which is important for the cellular NADP^+^/NADPH balance [25]) (Fig. 4A). Interestingly, the total folate levels in the parental and knockout parasites are not statistically different (Fig. 4D). This is likely due to the ability of apicomplexan parasites to uptake folates from the host cell via a high-affinity transporter [26, 27]. Interestingly, folate levels in the complemented cell line were lower than in the other strains, likely due to overexpression of DHFR-TS, which serves as a selectable marker in the complemented strain. While there are no detectable differences in intracellular folate or dihydrofolate levels among the strains, the total serine level is decreased in the knockout parasites (Fig. 4D), and this decrease can be reversed by LMF1 complementation. (Fig. 4D). Tetrahydrofolate, the product of DHFR-TS, is also decreased in the knockout cell line, consistent with the reduction in DHFR-TS activity. Complementation of LMF1 restored intracellular tetrahydrofolate levels to those of the parental strain. Downstream in the pathway, tetrahydrofolate is used by SHMT, a reversible enzyme that can form either serine or glycine. The intracellular serine levels are reduced in the knockout parasites but are restored in the complemented cell lines, suggesting that lowering tetrahydrofolate levels might influence subsequent pathways. Finally, we examined whether dTMP levels, the product of Thymidylate Synthase (TS) activity, were affected by the decrease in DHFR-TS activity. No statistically significant differences were found between parental and knockout cell lines (Fig. 4D).

To evaluate whether the decrease in DHFR-TS activity in the Δ*lmf1* parasites affected other substrates that can be influenced by pyrimidine metabolism, we monitored levels of glucose-6-phosphate (G6P), aspartate, and glutamine using untargeted metabolomics. G6P is directed into the Pentose Phosphate Pathway (PPP) to produce ribulose-5-phosphate. Aspartate is necessary for the synthesis of adenyl succinate and N-carbamoyl-aspartate, important intermediates for purine and pyrimidine synthesis, while glutamine provides nitrogen for pyrimidine nucleobases. Curiously, the Δ*lmf1* parasites exhibited lower intracellular levels of G6P and aspartate, a phenotype that is rescued in the complemented cell lines (Fig. 4D). However, total glutamine levels remained unchanged across all conditions analyzed. Although these lower levels of metabolites in the knockout cell lines may represent an increased activity of nucleotide synthesis, we cannot rule out the use of aspartate for energy metabolism and G6P to maintain the parasite’s redox balance and promote fatty acid synthesis. We measured total NADP^+^ and NADPH levels (Supplemental fig. S2) and observed a decrease in NADPH, correlating with reduced G6P, suggesting increased PPP use for redox balance and/or metabolite biosynthesis.

Previous studies have shown that folates can accumulate within the parasite’s mitochondrion [26]. Accordingly, we tested whether loss of LMF1 altered the uptake and distribution of folate within the parasite (Fig. 4E). For this purpose, we used the fluorescein-conjugated folate analog methotrexate (MTX), which is known to be transported into cells and mitochondria [26]. We incubated the parasites with MTX-fluorescein as previously described and observed the cells under a microscope. To visualize the mitochondrion, we co-stained the parasites with TMRE, a fluorescent indicator of mitochondrial membrane potential. There was a striking difference between the knockout and parental cell lines in relation to the pattern of accumulation of MTX within the cell (Fig. 4E). While the parental line showed a widespread distribution of the signal, we observed areas of accumulation that correspond to the mitochondrion (Fig. 4E). Meanwhile, the knockouts showed a stronger punctate signal, which does not colocalize with the mitochondrial signal (Fig. 4E). Since Δ*lmf1* parasites exhibit aberrant mitochondrial morphologies, we looked at MTX distribution in parasites showing a variety of previously defined mitochondrial morphologies (lasso, sperm, and collapsed) [6, 7]. Some mutant parasites exhibit concentrated staining within the mitochondrion and in areas resembling the nucleus (Fig. 4E, white arrows). We measured the total fluorescence of the MTX probe in the parasites and observed no difference between the parental and knockout strains, confirming the metabolomic results (Fig. 4F). We calculated the percentage of parasites exhibiting either mitochondrion-like or nucleus-like staining in both conditions. The mitochondrion-like staining is observed in all parental cell lines, whereas the knockout parasites present nuclear-like staining (Fig. 4G). These results indicate that, while folate uptake into the parasite is unaffected by the loss of LMF1, its accumulation within the mitochondrion is affected.

As folate accumulation across different cellular compartments was an unexpected result, we decided to examine the localization of DHFR-TS in parental and knockout parasites. For this purpose, we endogenously tagged DHFR-TS (TGGT1_249180) with a 3xHA tag in both strains and performed immunofluorescence using antibodies against the HA epitope tag and the mitochondrial marker TOM40. The parental cell line (ku80) showed diffuse nuclear staining, which colocalized with our DNA marker, DAPI (Fig. 4H). Additionally, there is a strong signal that colocalizes with TOM40 antibody staining, confirming that DHRF-TS may have a dual localization within the cell: the nucleus and the mitochondrion (Fig. 4H). Interestingly, the knockout parasites exhibit stronger nuclear HA-tag staining and lower mitochondrial protein levels, suggesting that a nuclear isoform predominates in the absence of LMF1 (Fig 4H). These results corroborate our previous observations of a distinct accumulation of folate in the parasites, likely due to the differential localization of DHFR-TS in the absence of LMF1.

### Coenzyme A metabolism is affected in the absence of LMF1

Another metabolism-related gene that appears to be downregulated upon LMF1 loss is Pank1. Pank1, along with Pank2, is part of the first step of Coenzyme A (CoA) biosynthesis. *Toxoplasma* can synthesize Coenzyme A from pantothenate, which can be imported into the parasite or synthesized from valine [28] (Fig. 5A). Valine is an essential amino acid for the parasite, which it must obtain from its host [29]. Once in the cytosol, valine is transported into the mitochondrion, where it is metabolized to pantoate. Pantoate is then transported back to the cytosol, where it is converted into pantothenate. Pantothenate is phosphorylated by either Pank1 or Pank2, initiating CoA biosynthesis (Fig. 5A). Pantothenate can also be taken up from the host. Interestingly, the only gene downregulated in the pathway is Pank1 (Fig. 5B). To confirm that the Pank activity was decreased, we measured the levels of enzymatic activity in crude extracts from intracellular parasites. The activity was indirectly measured by coupling Pank activity to pyruvate-kinase-lactate dehydrogenase, following the reduction of NADH at 340 nm (Fig. 5C) [30]. Δ*lmf1* parasites exhibit significantly lower activity compared to the parental cell line (Fig. 5C). Complementation of LMF1 reestablished normal enzymatic activity, indicating that the loss of LMF1 affects the pathway (Fig. 5C). Furthermore, we performed untargeted metabolomics to detect some of the metabolites related to CoA biosynthesis. Specifically, we monitored the intracellular levels of pantothenate, coenzyme A, and valine (Fig. 5D). Knockout parasites show higher levels of pantothenate when compared to the parental strain, indicating that this metabolite might be accumulating in the Δ*lmf1* parasites, likely due to the reduction of Pank activity (Fig. 5D). Interestingly, the levels of pantothenate remained elevated upon complementation (Fig. 5D). We also measured levels of CoA, the final product of this pathway. Intracellular levels of CoA are decreased in the Δ*lmf1* parasites, indicating that reducing Pank1 activity affects the formation of CoA. Normal levels of CoA were restored upon complementation. As explained above, valine transport from the host is essential for the parasite and must cross the plasma membrane and the mitochondrial membranes to be metabolized. We decided to check whether losing LMF1 would also affect intracellular valine levels. Δ*lmf1* parasites show lower levels of valine in comparison to both parental and complemented (Fig. 5D). Thus, our results confirm that LMF1, and likely the mitochondrion-IMC contact site, are important for the completion of CoA biosynthesis from pantothenate and/or valine.

**Figure 5.**
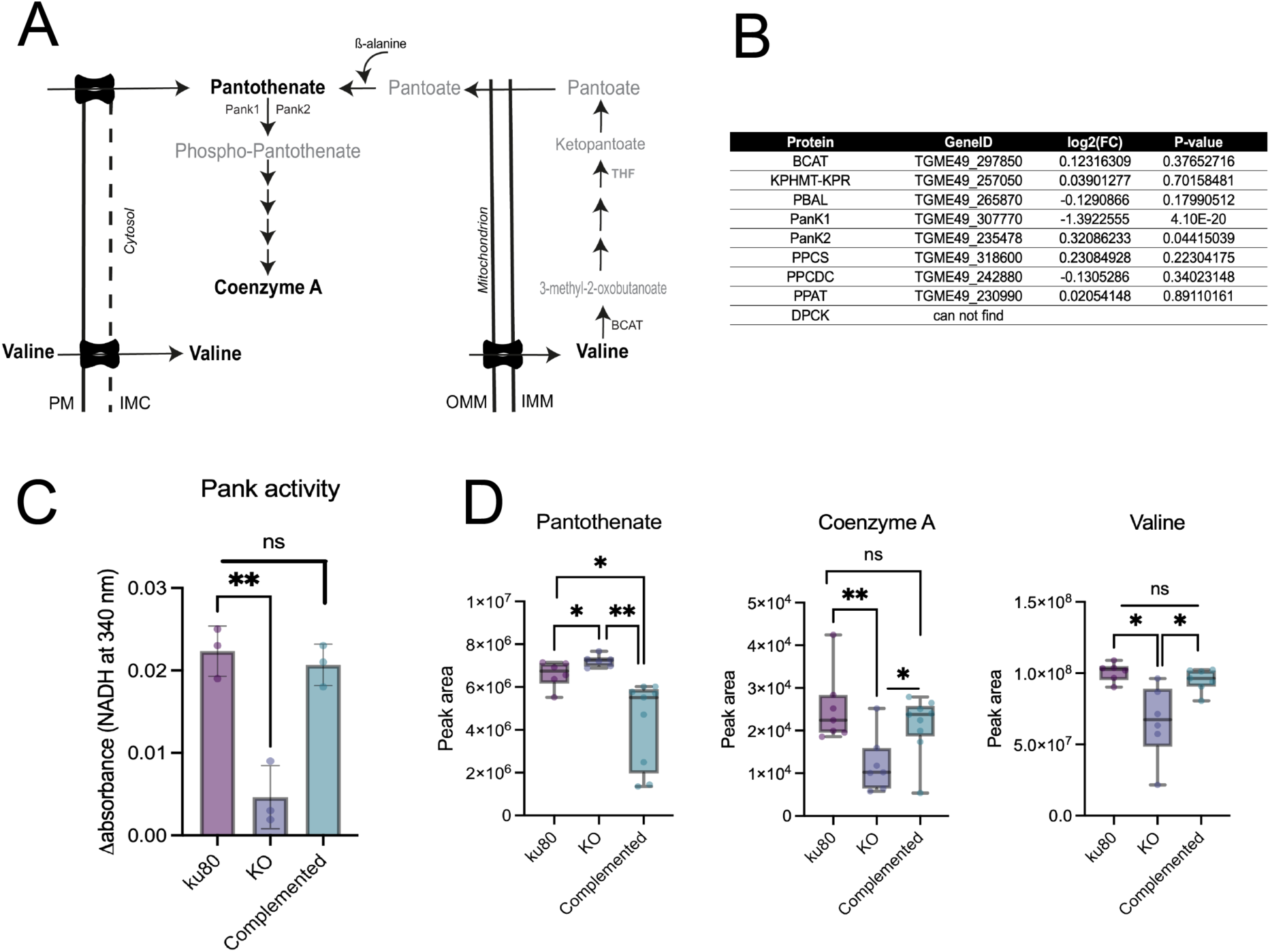
Loss of LMF1 impacts Coenzyme A biosynthesis. A) Schematic representation of the valine to CoA biosynthetic pathway. Adapted from [28]. PM = plasma membrane. IMC = inner membrane complex. OMM = outer mitochondrial membrane. IMM = inner mitochondrial membrane. BCAT = branched-chain amino acid transaminase. THF= tetrahydrofolate. Pank1/Pank2 = pantothenate kinase 1 and 2. B) Total activity of Pantothenate Kinase (PanK) in crude extracts of intracellular parasites. Enzyme activities were measured by observing the reduction of NAD^+^ into NADH at 340 nm. Enzymatic activity was performed in n=3. Data are means ± s.d. Statistical analysis is a Two-tailed unpaired t-test, ** P<0.01, ns = not significant. C) Total intracellular levels of valine, pantothenate, and coenzyme A. Metabolite quantification was performed in three technical replicates and three biological replicates (n=9). Data are means ± s.d.. Statistical analysis is a Two-tailed unpaired t-test, ** P<0.01; * P<0.05.

### Loss of LMF1 affects the parasite’s mitochondrial metabolism

Our results indicate a disruption in the metabolism of Δ*lmf1* parasites relative to the parental strain. A question that remains unanswered is whether mitochondrial activity is affected in the absence of LMF1. To assess the effect of LMF1 loss on mitochondrial activity, we measured the ability of mutant and parental parasites to consume oxygen under three different conditions: 1. glucose and glutamine as carbon sources (which mimics normal culture media); 2. glucose as the only carbon source, and 3. glutamine as the only carbon source (Fig. 6). A representative graph resulting from the assay is shown in Fig. 6A. The basal oxygen consumption rate (OCR) of the parental parasites is significantly higher than that of the Δ*lmf1* parasites in the presence of glucose/glutamine (230.4 ± 26.5 for the ku80 vs 213.5 ± 16 for the knockouts) or just glucose (184.2 ± 13.4 for the ku80 vs 140.6 ± 14.7 pmols of O_2_ for the knockouts) (Fig. 6B). Surprisingly, the knockout parasites exhibited higher oxygen consumption levels when provided with only glutamine as a carbon source (211.05 ± 16.7 vs 251.7 ± 27.6 pmols of O_2_ for the knockout) (Fig. 6A). This suggests a shift in preference regarding the substrates for oxidative phosphorylation in the knockout parasites. In addition to basal OCR levels, we measured glucose dependence by adding 2-deoxyglucose (2-DOG), an inhibitor of hexokinase, the first step in glycolysis. Corroborating our basal OCR observations, the parental cell lines showed more contribution of glucose by itself (decrease in 13.4 ± 6.17 for ku80 vs 3.47 ± 7.29 pmols of O_2_ for the knockout) or in combination with glutamine (decrease in OCR 33.9 ± 10.9 for the ku80 vs 19.9 ± 10.7 pmols of O_2_ for the knockout) (Fig. 6B). In the presence of just glutamine as a substrate, the parental parasites still showed a decrease in respiration when treated with 2-DOG (decrease in 2.03 ± 6.7829 pmols of O_2_), while the Δ*lmf1* parasites exhibited an increase in respiration in the presence of 2-DOG (2.92 ± 5.55 29 pmols of O_2_) (Fig. 6C). Next, we added the uncoupling agent FCCP, which dissipates the mitochondrial proton gradient and reveals the organelle’s maximal capacity. The parental cell line, when in the presence of glucose and glutamine, showed higher OCR values (94.2 ± 17.37 pmols of O_2_) in comparison to the knockout cell lines (106.3 ± 18.7 pmols of O_2_) (Fig. 6D). The same pattern, although less pronounced, was observed when glucose was the only substrate available, confirming a higher contribution of glucose to the metabolism of the parental cell lines. When incubated with glutamine as a sole source, the knockout cell line showed higher maximal respiration (167.9 ± 40.17 pmols of O_2_), almost 1.4X the increase in OCR presented by the parental line (121.2 ± 25.08 pmols of O_2_) (Fig. 6C). This confirms that Δ*lmf1* parasites preferentially use glutamine to fuel mitochondrial respiration.

**Figure 6.**
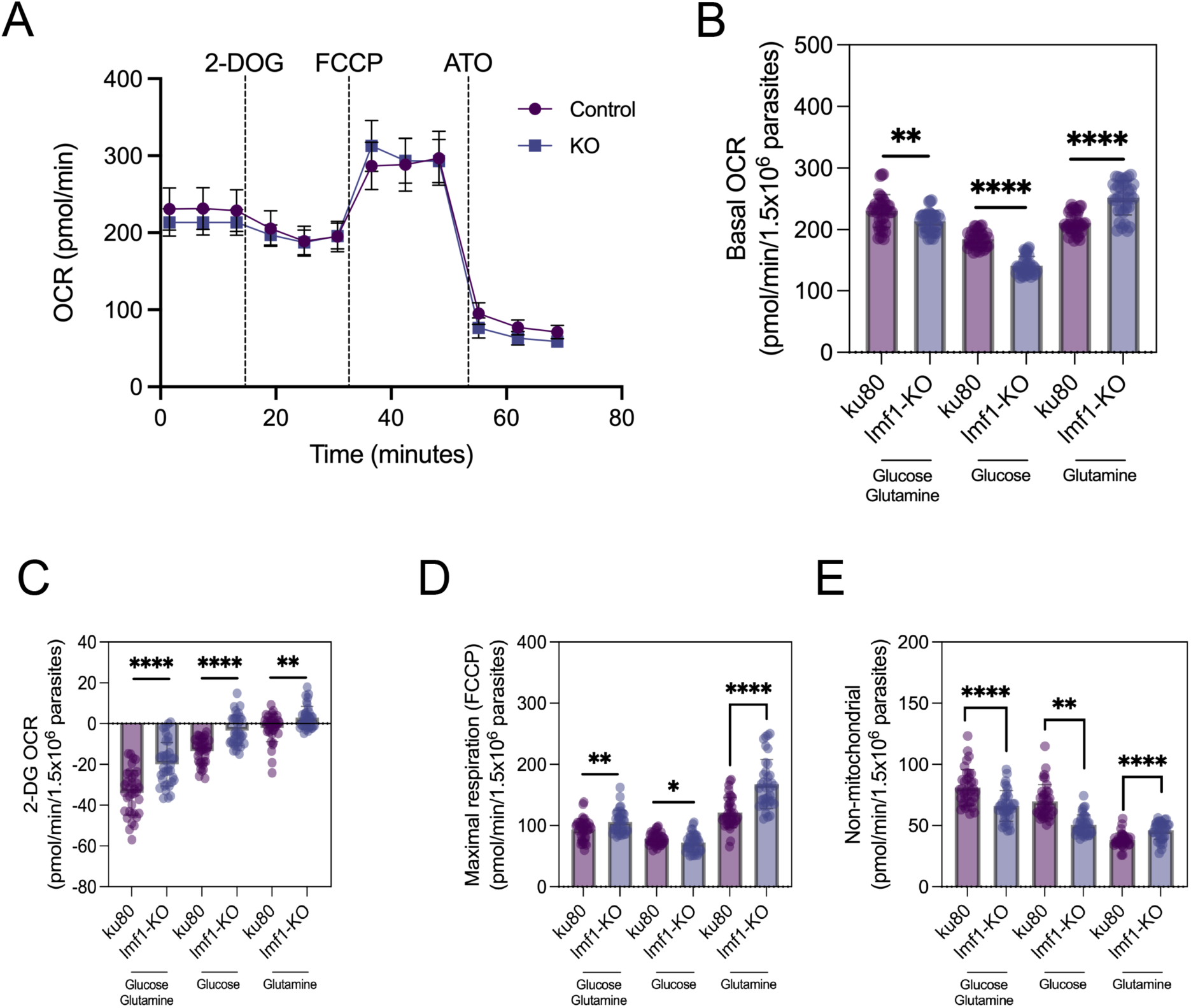
Glutamine is preferred over glucose in Δ*lmf1* parasites. A) Representative traces depicting OCR over time when supplying glucose and glutamine as ATP sources. B) Bar graphs representing the Basal OCR of parasites with different substrates. C) Bar graphs representing the OCR from parasites treated with 2-DOG. Note: the graph is shown in negative values to represent the decrease in OCR. D) Maximal respiration (FCCP) of parasites in different media conditions. D) OCR of non-mitochondrial respiration post-Atovaquone treatment. Data represent the mean ± SD of three technical replicates and are representative of three independent experiments. **** P<0.0001, *** P<0.001, ** P<0.01, * P<0.05 and ns not significant (P>0.05) (unpaired t-test, Two-tailed).

Finally, we inhibited mitochondrial respiration using Atovaquone, a well-characterized complex III inhibitor for *Toxoplasma*, and calculated the non-mitochondrial OCR (residual oxygen consumption utilized in other cellular processes) under all conditions (Fig. 6E). Parental cell lines showed a higher non-mitochondrial OCR after mitochondrial respiration inhibition in the presence or absence of glucose and glutamine. Moreover, higher non-mitochondrial OCR levels were observed for the knockout cell line in the presence of just glutamine. To confirm that Δ*lmf1* parasites do not prefer glucose as a main catabolic source, we performed plaque assays with the parental and knockout cell lines in media containing just glucose or no glucose (Supplemental fig. S2). Parental parasites can form plaques in the presence of just glucose, clearing more than 17% of the monolayer (17.3 ± 4.9%). In the absence of glucose, the parasites exhibited a slower growth phenotype, clearing 7.14 ± 3.3% of the monolayer. On the other hand, the knockout parasites already exhibited a reduction in plaque formation (4.8 ± 1.7%) when glucose was the sole available substrate. Without glucose, the differences in clearing are not statistically significant, confirming our previous results that glucose isn’t the preferred source for Δ*lmf1* parasites. In summary, Δ*lmf1* parasites exhibit a preference for utilizing glutamine as a carbon and ATP source, which is directly reflected in their cellular mitochondrial metabolism.

### Mitochondrial metabolism of Δ*lmf1* parasites resembles that of extracellular parasites

Our phenotypic characterization of the Δ*lmf1* parasites has shown that certain metabolic features, including mitochondrial respiration, are altered. We also observed differences in Ca^2+^-induced egress and microneme secretion, processes related to egress and invasion. One of the key differences between the parental and Δ*lmf1* strains is mitochondrial morphology, which in the knockout strain is collapsed, resembling that of extracellular parasites. Thus, it is plausible that the changes in metabolism and ion homeostasis in the knockout strain are related to the lack of LMF1 causing a shift towards an ‘extracellular state’. Accordingly, we analyzed whether intracellular and extracellular parasites differed in overall OCR. We compared the OCR of freshly syringe-released parasites (for more straightforward interpretation called as intracellular) to that of extracellular parasites that naturally egressed (parasites that completely lysed out of the host cells and stayed outside for a period of 12-16h, referred to as extracellular) in the parental and knockout cell lines in the presence of glucose and glutamine (Fig. 7). The parental line exhibited notable differences between intracellular and extracellular parasites parasites. The basal OCR (80.58 ± 12.39 for ku80 vs. 115.59 ± 5.56 pmols of O_2_ for the extracellular) was followed by a higher dependency on glucose for the intracellular parasites (decrease in 12.69 ± 6.79 vs. 29.47 ± 6.29 pmols of O_2_). Maximal respiration increased 2.3-fold in naturally egressed parasites compared with those released by syringe, indicating a higher mitochondrial maximal capacity. Interestingly, non-mitochondrial respiration was also higher in the extracellular parasites, which might indicate a high demand for other cellular processes (Fig. 7A). When analyzing the same parameters for the Δ*lmf1* parasites, no significant differences were observed in basal OCR, maximal respiration, or non-mitochondrial respiration between intracellular and extracellular parasites (Fig. 7B). Only a slight decrease in OCR was observed in extracellular parasites after treatment with 2-DOG (Fig. 7B). We performed a comparative analysis of the entire dataset using a heat map (Fig. 7C), which shows that the Δ*lmf1* parasites, both intra-and extracellular, are positioned intermediate to the parental parasites. To confirm that the syringe-lysed parental strain parasites, which we are considering ‘intracellular’, retained a lasso-shaped mitochondrion, we quantified mitochondrial morphology in both the syringed-lysed (intracellular) and naturally egressed (extracellular) parasites. As shown in (Fig. 7D), most of the parasites from the parental strain still presented a lasso-shaped mitochondrion after being syringe-released. Naturally egressed parental parasites, which have been extracellular for a prolonged period, exhibit the expected change in mitochondrial morphology. As expected, most mutant parasites show either collapsed or lasso-shaped mitochondria, whether syringe-lysed or naturally egressed. We conclude that Δ*lmf1* parasites exhibit mitochondrial respiration characteristics that fall between those of intracellular and extracellular parasites, likely due to defects in mitochondrial positioning.

**Figure 7.**
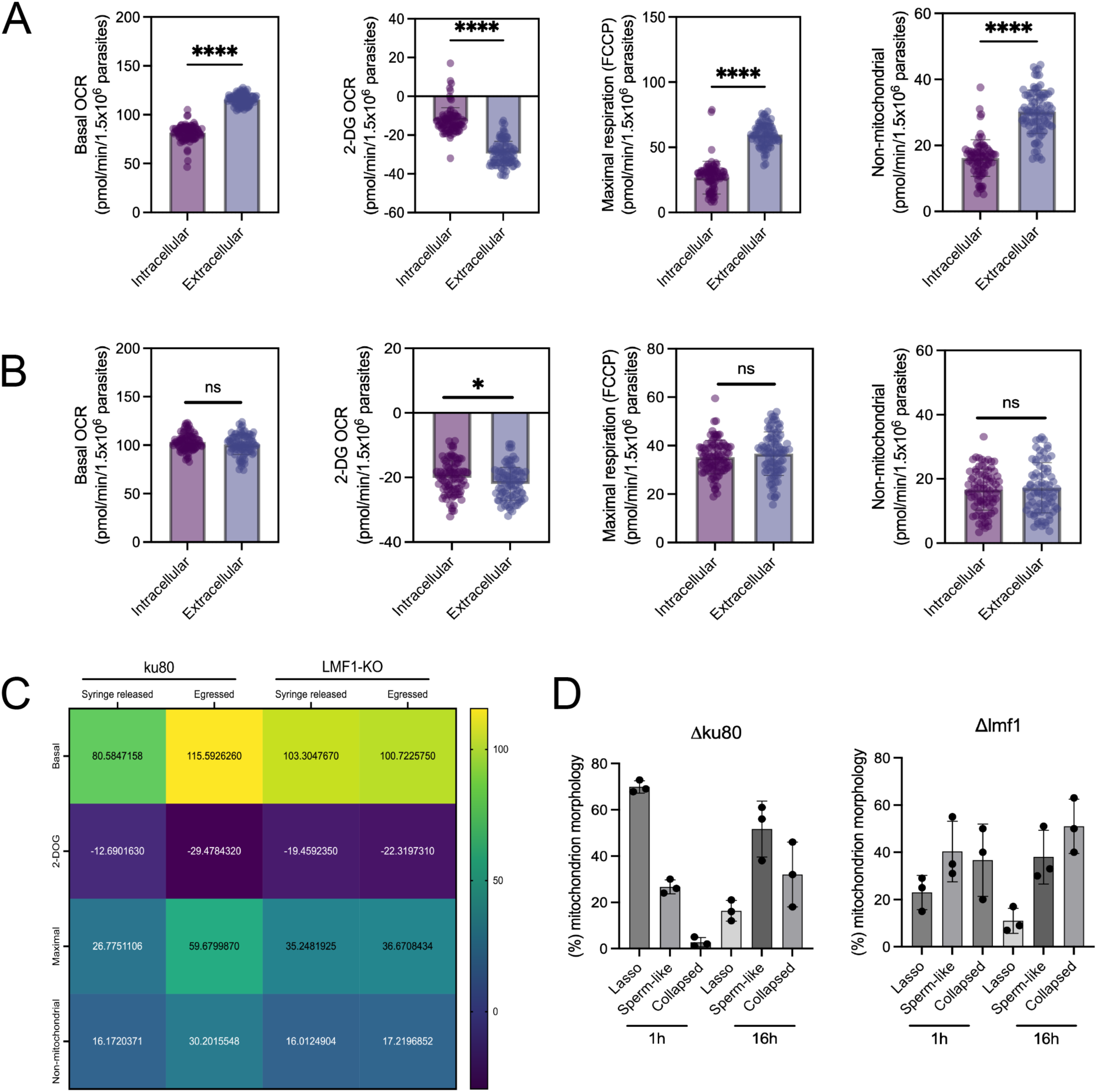
Δ*lmf1* parasites’ mitochondrial metabolism is similar to that of extracellular parasites. A) Bar graphs of Basal OCR, Glucose-dependent OCR (2-DOG), Maximal respiration, and non-mitochondrial respiration of intracellular and extracellular parasites of parental strain (**Δ**ku80). B) Bar graphs of Basal OCR, Glucose-dependent OCR (2-DOG), Maximal respiration, and non-mitochondrial respiration of intracellular and extracellular of LMF1 knockout parasites (**Δ***lmf1*). C) Heat-map showing the average values of each parameter evaluated in this assay, warm colors represent the higher values, cold colors represent the lower values. D) Mitochondrial morphology counts of parasites from both strains, 1h post-release from the host cells (in this work, called intracellular parasites), and 16h after complete egress (extracellular parasites). Data represent the mean ± SD of three technical replicates and are representative of three independent experiments. **** P<0.0001, *** P<0.001, ** P<0.01, * P<0.05 and ns not significant (P>0.05) (unpaired t-test, Two-tailed).

## DISCUSSION

Membrane contact sites (MCSs) are areas of close apposition between organelles, allowing coordination of their activities [31]. These contact sites are important for many biological processes, including lipid exchange [32] and ion homeostasis [33]. Given these key roles, MCSs are a focus of research in many pathogenic eukaryotes, including *Toxoplasma* [34, 35]. It has been reported that there are zones of membrane proximity between the mitochondrion and apicoplast [9], the apicoplast and ER [36], the mitochondrion and IMC [6], the mitochondrion and ER [37], and the mitochondrion and nucleus [38]. So far, we know that an apicoplast Two Pore Channel (TPC) mediates contact between the apicoplast and the ER, which is important for Ca^2+^ transfer between the organelles [39]. The parasite’s mitochondrion can interact with the ER via VDAC, and loss of this contact site leads to mitochondrial morphological abnormalities and defects in purine metabolism [37]. The mitochondrion also interacts with the nucleus via two pore-forming proteins, Nup503 (located in the nucleus) and ATOM40 (located in the mitochondrion). Knockdown of Nup503 leads to aberrant mitochondrial morphology and loss of MCS with the nucleus [38]. While progress has been made in identifying the proteins involved in forming the contact sites, the functions of the association between membranes and organelles remain to be defined. Building on the identification of the drivers of tethering between the mitochondrion and the IMC in *Toxoplasma*, we have begun characterizing the functional roles of this contact site.

One of the canonical roles of a contact site is the transfer of metabolites and ions among organelles. Our results suggest that contact between the mitochondrion outer membrane and the IMC in *Toxoplasma* is important for the metabolism of important cofactors, such as tetrahydrofolate and Coenzyme A. Interestingly, we showed that loss of LMF1 and, consequently, the contact between the mitochondrion and the IMC, results in downregulation of DHFR-TS. In mammalian cells, folate metabolism is compartmentalized between the mitochondria and the cytosol [40], and DHFR is located in both compartments [41]. Thereby, mitochondrial folate metabolism contributes to one-carbon metabolism, whereas the cytosolic pathway contributes to purine metabolism [42]. Protozoan parasites such as *Toxoplasma*, *Plasmodium*, *Cryptosporidium*, and *Leishmania* possess an unusual bifunctional DHFR-TS [12]. Recent studies have revealed a sophisticated host response mechanism to *Toxoplasma* infection that restricts folate availability, confirming the importance of this metabolite for parasite survival [15]. The same study overexpressed the parasite’s DHFR-TS and localized it to the nucleus [15], which aligns with the results of whole-cell subcellular proteomics [14]. Given our observation that disruption of mitochondrial morphology affects the expression of this enzyme, we revisited its localization. Our data showed that an endogenous DHFR-TS tag results in dual localization: the nucleus, as previously reported, and the mitochondrion. LMF1 knockout parasites have a stronger staining for the tagged DHFR-TS in the nucleus. More studies are necessary to understand the functional importance of the localization pattern observed for DHFR-TS.

Previous studies have shown that a fluorescent folate analog accumulates in mitochondria [26]. Using a fluorescent folate probe, we observed that Δ*lmf1* parasites still take up folate. However, mitochondrial transport may be affected, suggesting that the mitochondrial-IMC tether mediates mitochondrial folate uptake. More detailed metabolomics is needed to understand the role of the contact site in folate metabolism. Interestingly, knockdown of VDAC in *Toxoplasma* leads to the accumulation of certain pyrimidine intermediates, suggesting that mitochondria play a critical role in the pyrimidine salvage pathway [37]. We observed decreased intracellular levels of G6P, aspartate, and serine. Although we don’t have a clear explanation for the decrease in G6P and aspartate, these lower levels might indicate that either serine uptake from the host is affected or that the parasites are synthesizing more lipids, such as phosphatidylserine. Metabolite levels were restored by complementing the knockouts with *LMF1*, indicating that this protein is necessary to maintain the intracellular balance of these metabolites. More detailed metabolomics and lipidomic studies are needed to understand the parasites’ metabolic remodeling when they are unable to form proper contact sites.

Another metabolic enzyme downregulated in Δ*lmf1* parasites is Pank1. Pank1 phosphorylates pantoate in the cytoplasm, initiating CoA biosynthesis. The coenzyme A (CoA) biosynthesis pathway is essential for *Toxoplasma* survival in vitro [28, 43] and for cyst formation [28], and has been proposed as a promising drug target against *Toxoplasma* and *Plasmodium* [44]. However, pantothenate can also be transported from the host and converted into CoA using the cytosolic pathway. Our observation that *Pank1* is downregulated in the *Δlmf1* parasites supports the hypothesis that the contact site mediated by LMF1 is critical for metabolic activities. *Toxoplasma* shows incredible metabolic flexibility in culture [45]. It has been shown that cooperation between glucose and glutamine metabolism is essential for the parasite’s lytic cycle in vitro [46]. When we compare basal oxygen consumption rates between knockouts and parental parasites, we observe a significant decrease in mitochondrial activity in the knockout parasites in the presence of glucose. Remarkably, *Δlmf1* parasites use glutamine more effectively than glucose to boost mitochondrial respiration. Previous work has shown that extracellular parasites accumulate higher levels of glucose intermediates, but it is unclear whether this is due to increased glucose consumption or to a slower downstream pathway for glucose degradation [47]. Our data suggest that differences in mitochondrial morphology may correlate with a metabolic preference for glutamine as the primary ATP-producing substrate in extracellular parasites. Glutamine metabolism is active in both intracellular and extracellular parasites, contributing to the TCA cycle metabolite pool in both forms [46]. Extracellular tachyzoites can store high amounts of gamma-aminobutyric acid (GABA), and this metabolite can be used to fulfill mitochondrial metabolism [47]. As we haven’t measured the total GABA levels in these parasites, we can’t confirm whether this pathway is upregulated upon LMF1 loss.

Egress is a critical step in the parasite’s lytic cycle, marking the transition from the intracellular to the extracellular environment. Different stimuli and second messengers, such as K^+^, Ca^2+^, and cyclic nucleotides, can regulate natural egress [21, 48]. We were interested in DGK3 over other genes that appeared to be upregulated because of the mitochondrial morphology changes that accompany egress [6]. DGKs convert diacylglycerol to phosphatidic acid as part of the phosphoinositide-phospholipase C signaling cascade [21]. In *Toxoplasma*, this triggers the release of Ca^2+^ from intracellular stores and the secretion of micronemes, leading to parasite egress. *Toxoplasma* has three DGKs; DGK1 is essential, while DGK2 and 3 are dispensable [18]. DGK3 is localized to the micronemes, and while it is dispensable, it still impacts parasite propagation by contributing to Ca^2+^ release and microneme secretion [19]. The upregulation of DGK3 in the mutant strain might indicate that mitochondrial morphology and function influence egress signaling. The knockout parasites are more sensitive to Ca^2+^-induced egress, and our experiments have shown that they exhibit higher intracellular Ca^2+^ levels and differences in microneme positioning. It remains unclear what role *Toxoplasma*’s mitochondrion plays in Ca^2+^ signaling, as key proteins required for this process are absent from the parasite’s genome, and VDAC depletion did not alter the parasite’s Ca^2+^ levels [37]. We speculate that this phenotype may be due to differences in the contact sites between the IMC and the ER. During egress, some Ca^2+^ is released from the ER and distributed to other cellular compartments, likely via contact sites [49]. Egress and invasion events are tightly regulated by Ca^2+^ fluctuations within the parasite [50, 51]. As the extracellular medium contains more calcium than the intracellular medium [52], we hypothesize that *Δlmf1* intracellular parasites behave more like extracellular parasites. More work is needed to uncover the signaling pathway linking egress and mitochondrial morphological changes.

Our data indicate that mitochondrial positioning is a key player in the parasite’s environmental sensing, leading to changes in metabolic and intracellular signaling. We posit that intracellular parasites with incorrect mitochondrial positioning appear be in a pseudo-extracellular state or to have adapted to facilitate efficient egress and invasion. In summary, our work shows for the first time that the mitochondrion and the pellicle form a contact site that mediates metabolite uptake and facilitates egress signaling.

## METHODS

### Parasites and host cells

All the parasite strains were maintained via continued passage through human foreskin fibroblasts (HFFs) purchased from ATCC (SCRC-1041) and cultured in Dulbecco’s modified Eagle’s medium (DMEM) high glucose, supplemented with 10% fetal calf serum (FCS), two mM L-glutamine, and 100 U penicillin/100 µg streptomycin per ml. The cultures were maintained at 37°C and 5% CO_2_. Parental parasites and derivatives used in this study were of the strain RH lacking hypoxanthine-xanthine-guanine phosphoribosyl transferase (HPT) and Ku80 (RHΔ*hpt*Δ*ku80*) [53]. Cells were inspected for mycoplasma contamination using a PCR-based VenorGeM Mycoplasma Detection Kit (Sigma, MP0025-1KT).

### RNA sequencing

Parental and Δ*lmf1* parasites were grown for 18 hours in host cells. Parasites were harvested with host cells by scraping in cold PBS, followed by centrifugation at 2000 × g for 5 min at 4°C. The pellet was resuspended in 10 mL PBS using a syringe to release parasites from host cells, and the samples were centrifuged again. To extract the RNA, the pellet was treated with 1 mL of TRIzol for 5 minutes at room temperature, then with 200 μL of chloroform, and centrifuged at 12,000 × g for 15 minutes at 4°C. The aqueous phase was treated with 500 μL of chloroform again and centrifuged. The aqueous phase was mixed with 500 μL of isopropanol, incubated at room temperature for 10 minutes, and centrifuged at 12,000 × g for 10 minutes. The RNA was washed with 1 mL of 75% ethanol and centrifuged at 7,500 × g for 5 minutes. The resulting RNA pellet was air-dried and resuspended in 50 μL of nuclease-free water. Each condition was performed in biological triplicate. Samples were stored at −80°C prior to shipment to AZENTA for library construction and Illumina Next-Generation Sequencing. For each sample, ∼30 M 2x150 bp pair-end reads were obtained. Analysis of the raw RNA sequencing data was performed using the Galaxy online platform. First, FastQC was used to confirm the quality of the data. Trim Galore was then used for further quality control and adaptor trimming. The data were then mapped to the *Toxoplasma* genome using hiSAT2, and subsequently, ht-seq-count was used to count the number of reads that aligned to each gene. Finally, DESEQ2 was used to identify individual genes from the RNA sequencing data. Genes of significance were identified by an adjusted p-value <0.05 and a log2 score >1 or <- 1. Genes were identified using ToxoDB.org [54].

### Phosphatidic acid uptake assay

The protocol was performed as previously published by Katris with some modifications. Briefly, parasites were syringe-released, filtered, and harvested by centrifugation at 1,000 × g for 10 min. Parasites were washed once in complete DMEM and resuspended in DMEM supplemented with 10 mM HEPES to a final concentration of 5x10^7^ parasites. Parasites were then transferred to pre-warmed DMEM without FBS, supplemented with 10 mM HEPES and 10 µM NBD-PA (Avanti Research). Parasites were incubated in this solution for 30 seconds. The reaction was stopped by placing the parasites on ice for 2 minutes. After quenching, the samples were centrifuged (8,000 × g for 5 min) and washed once with PBS. After that, parasites were fixed with 4% paraformaldehyde and placed into coverslips coated with poly-D-lysine. Samples were blocked in 2% FBS for 1 h and stained with an anti-SAG1 antibody (1:5,000) for 1 h at room temperature. PA uptake was quantified using ImageJ, measured in the FITC channel. The ePA/SAG1 ratio was calculated by dividing the fluorescence intensity in channel 1 by that in channel 2.

### Egress assay

The protocol was performed as described in [55]. HFFs were seeded into 94-well plates at least 72h before infection. Approximately 24 h before the experiment, the host cells were infected with 5×10^^4^ parasites per well. A row of wells was left uninfected as a control. Infected cells were washed once with PBS, and the media were replaced with HBSS only (control), 1 µM A23187 only, or 30 µM R50922. For the ionophore sensitivity assay, the parasites were treated with different concentrations of A23187 (0.1, 0.25, 0.5, and 1μM). Plates were placed in a water bath at 37 °C for 5 minuntes. Egress was halted by centrifuging the plates at 500 x g for 5 min at 4°C, to pellet egressed parasites and cell debris. Carefully, 50 μl of the supernatant was transferred to a new plate. The uninfected wells were then lysed by adding 10 μl of the lysis solution provided with the Cytotox Assay kit, scraping the monolayer with a pipette tip, and pipetting up and down to resuspend the cells. 50 μl of the resulting cell lysate was transferred to the new plate. Quantification of egress was performed by using the CytoTox Cell Kit (Promega) as recommended by the manufacturer. Absorbance readings were obtained at 490 nm, using a BioTek Synergy H1 Multimode Reader spectrophotometer.

### Calcium measurements

Parasite calcium concentration was determined as previously described in [56, 57] with some modifications. Briefly, intracellular parasites were syringe-released, filtered, washed, and then centrifuged twice at 500 × g for 10 minutes at room temperature in buffer A (BAG, 116 mM NaCl, 5.4 mM KCl, 0.8 mM MgSO4, 5.5 mM D-glucose, and 50 mM HEPES, pH 7.4). Then, parasites were resuspended to a final density of 2×10^7^ parasites/ml in Ringer buffer (155 mM NaCl, 3 mM KCl, 1 mM MgCl_2_, 3 mM NaH_2_PO_4 ·_ H_2_O, 10 mM HEPES, pH 7.3-, and 5 mM glucose; plus 1.5% sucrose and 5μM of Fura 2-AM). The suspension was incubated for 30 minutes at 28°C in a water bath with mild agitation. Subsequently, the parasites were washed twice with Ringer’s buffer to remove extracellular dye. Parasites were resuspended to a final density of 2×10^7^ parasites/ml in Ringer’s buffer and kept on ice. For fluorescence measurements, 2×10^6^ parasites/well were added to a UV-sensitive 96-well plate. The plate was placed in a thermostatically controlled BioTek Synergy H1 Multimode Reader spectrophotometer under conditions for Fura-2-AM excitation (340 and 380nm) and emission (510nm). The Fura-2-AM fluorescence response to intracellular Ca^2+^ concentration ([Ca^2+^] _i_) was calibrated from the 340/380 nm ratio after subtraction of the background fluorescence of the cells at 340 and 380 nm.

### Immunofluorescence assays

For all immunofluorescence assays (IFAs), infected HFF monolayers were fixed with 3.5% paraformaldehyde, quenched with 100 mM glycine, and blocked with PBS containing 3% bovine albumin serum (BSA). Cells were permeabilized in PBS containing 3% BSA and 0.25% Triton X-100 (TX-100). Samples were then incubated with primary antibodies diluted in permeabilization solution for 1 h, washed five times with PBS, and incubated with the respective Alexa Fluor-conjugated antibodies and 5 µg/ml Hoechst 33342 (Thermo Fisher Scientific; cat. no. H3570) in PBS for 1 hour. The coverslips were washed five times with PBS. After washing, the coverslips were mounted in ProLong Diamond (Thermo Fisher Scientific). Image acquisition and processing were performed using either a Nikon Eclipse 80i microscope with NIS-Elements AR 3.0 (objective lens 100×/1.40 Oil Plan Apochromat, z-step size of 0.3 µm), or a Zeiss LSM 800 AxioObserver microscope with an AiryScan detector using a ZEN Blue software (version 3.1). The images in this microscope were acquired using a 63x Plan-Apochromat (NA 1.4) objective lens. All images were acquired as Z-stacks with an XY pixel size of 0.035 µm and a Z-step size of 0.15 µm. All images were then processed with Airyscan using ZEN Blue (Version 3.1, Zeiss, Oberkochen, Germany). Images were processed and analyzed using FIJI ImageJ 64 Software. Primary antibodies used in this study are rabbit anti-HA (C29F4 Cell Signaling, 1:1000 dilution), rabbit anti-ATOM40 [58] (1:1000 dilution), and rabbit anti-MIC5 [59] (1:1000 dilution). Secondary antibodies included Alexa Fluor 594- or Alexa Fluor 488-conjugated goat anti-rabbit-IgG and goat anti-mouse-IgG (Invitrogen), all used at 1:1000.

### Microneme secretion

Freshly lysed parasites were harvested, filter-purified, and centrifuged at 1000 x g for 10 minutes. Parasites were resuspended in invasion medium (DMEM/20 mM HEPES/3% FBS) at a final concentration of 10^9^ parasites/ml. Aliquots of 100 of the parasite suspension were used for the secretion assay for each sample, one for ethanol stimulation, and the other for natural secretion without stimulation. For ethanol stimulation, 2% ethanol was added. For drug-stimulated microneme secretion, 2 µM of A23187 and 30 µM of R50922 were added. Samples were incubated at 37 °C for 10 minutes, and the reaction was stopped by halting secretion after the parasites were incubated on ice for 5 minutes. Samples were centrifuged at 1000 × g for 10 minutes at 4°C. After centrifugation, 80 µl of supernatant and the pellet were collected. The supernatant was centrifuged again, and 60 µl of the new supernatant was used for Western blotting. Pellets were lysed with RIPA buffer and prepared for western blotting as described above.

### Western blot

Protein samples were resuspended in 2x Laemmli sample buffer (Bio-Rad) with 5% 2-mercaptoethanol (Sigma-Aldrich). Samples were boiled for 5 min at 95°C before separation on a gradient 4–20% SDS-PAGE gel (Bio-Rad). Samples were then transferred to a nitrocellulose membrane using standard semidry transfer protocols (Bio-Rad). Membranes were probed with. Membranes were then washed and probed with either goat anti-mouse-IgG horseradish peroxidase or goat anti-rabbit-IgG horseradish peroxidase (Sigma-Aldrich) at a dilution of 1:10,000 for 1 h (GE Healthcare). Proteins were detected with the SuperSignal West Femto substrate (Thermo Fisher) and imaged with the FluorChem R system (Biotechne). Full original Western blots are shown in the Supplemental fig. S2.

### DHFR-TS activity assay

Freshly egressed parasites were harvested and washed twice by centrifugation (1,000 x g for 10 minuntes) in PBS. Pellets were then resuspended in Cell Lysis Buffer (Invitrogen) containing Protease Inhibitor Cocktail (Thermo) and subjected to freeze-thaw cycles in liquid nitrogen. Samples were centrifuged at 11,000 x g for 30 minutes, and the resulting supernatant was collected. Total protein quantification was performed using the BCA assay kit (Pierce). A total of 100 μg of total extract was used per enzymatic assay. The DHFR-TS assay was performed using the DHFR Assay Kit (Sigma) according to the manufacturer’s instructions in a UV-sensitive 96-well plate. The plate was placed in a thermostatically controlled BioTek Synergy H1 Multimode Reader spectrophotometer.

### Endogenous tagging of DHFR-TS

For the C-terminal endogenous tagging of *DHFR-TS*, we introduced a cassette encoding a 3x-HA tag directly upstream of the stop codon. This cassette included the selectable marker HXGPRT and was amplified from the vector pLIC-3xHA-DHFR [53] with primers that included the homology regions of each gene to promote recombination. The cassette insertion was facilitated by CRISPR. For this purpose, we replaced the guide RNA in pSAG1-Cas9-GFP-pU6-sgUPRT [60] for one targeting the gene of interest (GOI) locus using the Q5 site-directed mutagenesis kit (NEB). Primers used include those for the sgRNA (Fw: actcttcatggttttagagctagaaatagcaag; Rv: catcccttttaacttgacatccccatttac) and for the tagging cassette (Fw: gctacgtcccgcacggacgaatccagatggagatggctgtcaattaaaattggaagtggagg; Rv: gttttcccagtcacgacgaagaaaacaaggcgaggtgagactgtgtgaaatgccacat). Freshly egressed parasites were harvested, washed once in PBS, resuspended in transfection buffer (Buffer P3, Lonza), and transferred to transfection cuvettes. A total of 2×10^7^ parasites of the RHΔ*ku80* [53] cell line or the RHΔ*ku80*Δ*lmf1* [7] were used in each transfection, with 1 µg of the cassette and 1 µg of the Cas9 plasmid, using the Lonza nucleofection system [10]. Parasites were selected using pyrimethamine, and independent clones were collected by serial dilution.

### Metabolite extraction and metabolomic analyses

Samples were stored in -80°C until the day of extraction and instrument analysis. All samples were extracted with ice-cold 80% methanol. The samples were spiked with isotopically labeled internal standards. The internal standards were 500 ng of d3-Acetyl-CoA, 50 ng of 13C5-N-Folic acid, and 50 ng of 13C3-Serine. Samples were extracted by sonication in a chilled water bath at 4°C, vortexed, then centrifuged at 10,000 g for 10 minutes at 4°C. The extracted supernatants were collected and dried using nitrogen gas. For analysis, the samples were reconstituted in 0.1 mL of 50% acetonitrile. An Agilent 1290 Infinity II liquid chromatography (LC) system, coupled to an Agilent 6470 series QQQ mass spectrometer (MS/MS), was used to analyze the samples. (Agilent Technologies, Santa Clara, CA). To achieve better coverage, two types of liquid chromatography were used to analyze each sample. For reverse phase analysis, a Waters Atlantis T3 2.1 mm x 150 mm, 3.0 µm column was used for LC separation (Waters Corporation, Milford, MA). The buffers were A) water + 0.1% formic acid and B) acetonitrile + 0.1% formic acid. The linear LC gradient was as follows: time 0 minutes, 0 % B; time 1 minute, 0 % B; time 18 minutes, 100 % B; time 20 minutes, 100 % B; time 20.5 minutes, 0 % B; time 25 minutes, 0 % B. The flow rate was 0.3 mL/min, and the column was heated to 30°C. For hydrophilic interaction chromatography (HILIC) analysis, an Imtakt Intrada Amino Acid column 2.0 x 150 mm, 3.0 µM was used (Imtakt USA, Portland, OR). The buffers were A) acetonitrile + 0.3% formic acid and B) 20% acetonitrile in water containing 80% 100 mM ammonium formate. The linear LC gradient was as follows: time 0 minutes, 0 % B; time 1 minute, 0 % B; time 18 minutes, 100 % B; time 20 minutes, 100 % B; time 20.5 minutes, 0 % B; time 25 minutes, 0 % B. The flow rate was 0.3 mL/min, and the column was heated to 30°C. Multiple reaction monitoring was used for MS analysis. Data were acquired in both positive and negative electrospray ionization (ESI) modes. The jet stream ESI interface had a gas temperature of 325°C, a gas flow rate of 9 L/minute, a nebulizer pressure of 35 psi, a sheath gas temperature of 250°C, a sheath gas flow rate of 7 L/minute, a capillary voltage of 3800 V in positive mode and 3500 V in negative mode, and a nozzle voltage of 1000 V. The ΔEMV voltage was 500 V. Agilent MassHunter Quantitative Analysis software (version 10.1) was used for data analysis.

### Folate uptake assay

The protocol was performed as previously described [26], with a few modifications. Fluorescein methotrexate (Molecular Probes) was resuspended to a concentration of 1 mM in alkaline water. Intracellular parasites were syringe-released, filtered, washed twice in PBS, and resuspended in PBS to a final concentration of 5 × 10^6^ parasites per sample. Parasites were incubated with 10 µM fluorescent MTX for 30 minutes at 37 °C. In the last 10 minutes, we added 25 nM of Tetramethylrhodamine ethyl ester (TMRE). Parasites were washed twice in PBS to remove excess reagents. Images were acquired with a Zeiss LSM 800 AxioObserver microscope equipped with an Airyscan detector and ZEN Blue software (version 3.1). The images in this microscope were acquired using a 63× Plan-Apochromat (NA 1.4) objective lens. All images acquired from this microscope were acquired as *Z*-stacks with an XY pixel size of 0.035µm and a *Z*-step size of 0.15 µm. All images were then processed with Airyscan using ZEN Blue (Version 3.1, Zeiss, Oberkochen, Germany). Images were processed and analyzed using FIJI ImageJ 64 Software.

### PanK activity

The activity of PanK was indirectly measured in crude extracts as previously described [61], with the following modifications: intracellular parasites were harvested by syringe release using a 27-gauge needle, filtered, and then centrifuged at 2,000 × g for 5 min at 4°C. After 3 washes in PBS, the parasite pellet was resuspended in parasite lysis buffer (50 mM Tris-HCl, 150 mM NaCl, 0.5% NP-40, pH=7.4). Parasites were homogenized by 3 cycles of freeze-thaw. The supernatant was recovered after centrifugation at 12,000 × g for 30 min at 4°C. Reaction buffer contained 1mM phosphoenolpyruvate (PEP), 33U of PK/LDH (Sigma), 0.2 mM NADH, 10 mM MgCl_2_, 100 mM Tris-HCl, 2 mM ATP, 5 mM Pantothenate, and 100 μg of parasite extract, in a final volume of 150 μL. Reaction was started by the addition of Pantothenate. All the assays were performed in a UV-sensitive 96-well plate. The plate was placed in a thermostatically controlled BioTek Synergy H1 Multimode Reader spectrophotometer for analysis.

### Respiration assays

The basal oxygen consumption rate (OCR) and extracellular acidification rate (ECAR) were measured using a Seahorse XF 96 Pro Analyzer. The experimental protocol was conducted as previously described [62, 63]. Briefly, parasites were harvested and washed twice in XF Base Media (Agilent) at pH 7.4. After washing, the parasites were resuspended in Base Media supplemented with the corresponding substrate (5 mM glucose, 2 mM glutamine, or a combination of both), and 1.5 x 10^6^ parasites were seeded onto the pre-coated 96-well plates. To evaluate maximal respiration, carbonyl cyanide-p-trifluoromethoxyphenylhydrazone (FCCP) was added to a final concentration of 1 µM. Atovaquone was added to a final concentration of 1 µM to inhibit the electron transfer chain (ETC).

### Statistical analyses

Data were analyzed using GraphPad Prism software (version 9.00, La Jolla, CA). Analyses were performed using either an unpaired, one-tailed Student’s *t*-test or a one-way ANOVA with Tukey post-correction.

## Supplemental Figures

**Supplemental Figure S1.**
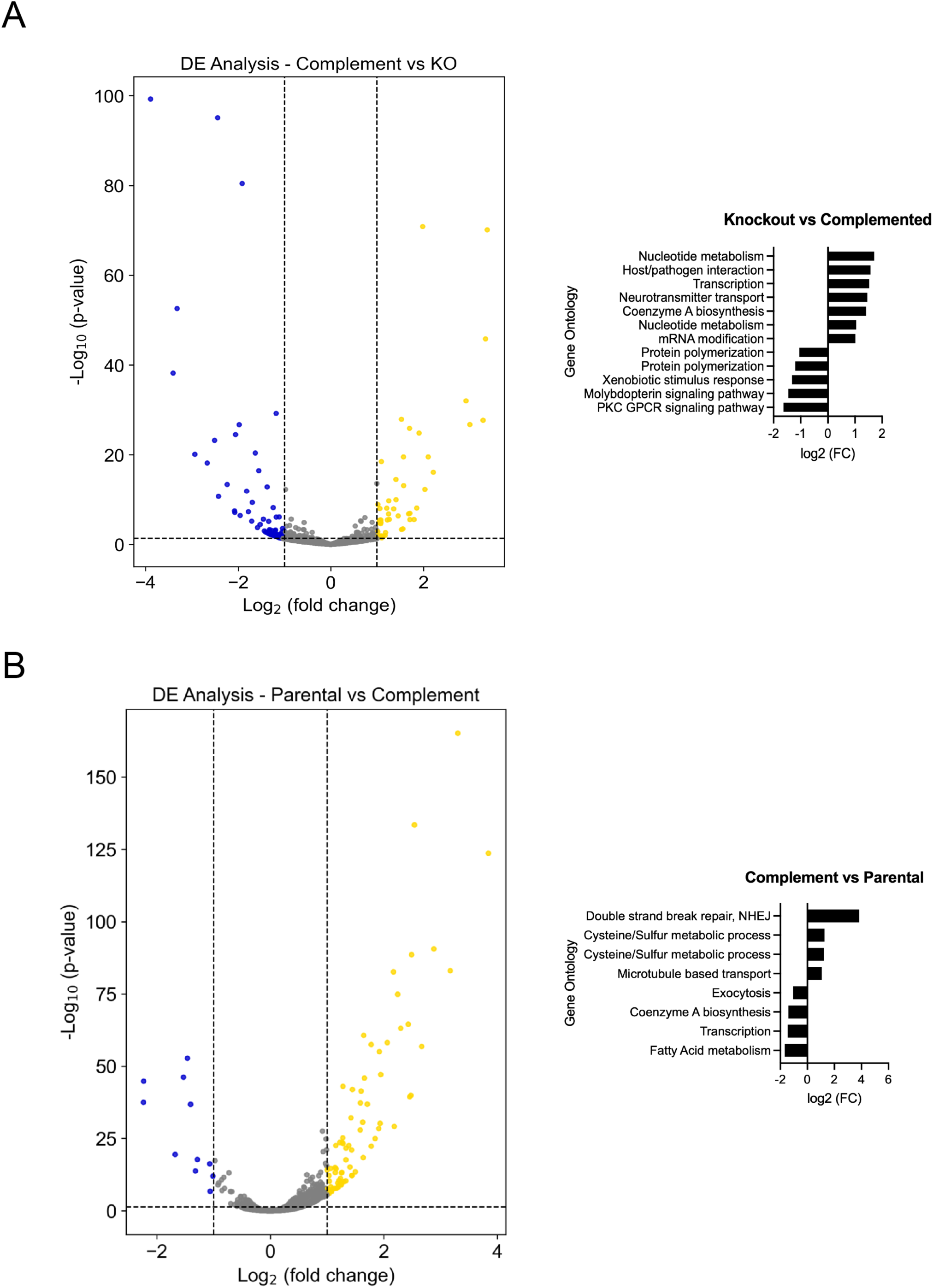
A) Volcano plot showing all differentially expressed genes with a log2 Fold Change >0.5 and p < 0.05 in complemented (Δ*lmf1*+*LMF1-HA)* parasites in comparison to the Δ*lmf1* strain. Genes of interest are highlighted with colored circles: yellow = upregulated genes of interest; blue = downregulated genes of interest. Bar graph showing the Gene Ontology (GO) for the main pathways up- and downregulated in the strain. B) Volcano plot showing all differentially expressed genes with a log2 Fold Change >0.5 and p < 0.05 in complemented (Δ*lmf1*+*LMF1-HA)* parasites in comparison to the parental strain. Genes of interest are highlighted with colored circles: yellow = upregulated genes of interest; blue = downregulated genes of interest. The experiment was performed in triplicate. Bar graph showing the Gene Ontology (GO) for the main pathways up- and downregulated in the strain.

**Supplemental Figure S2.**
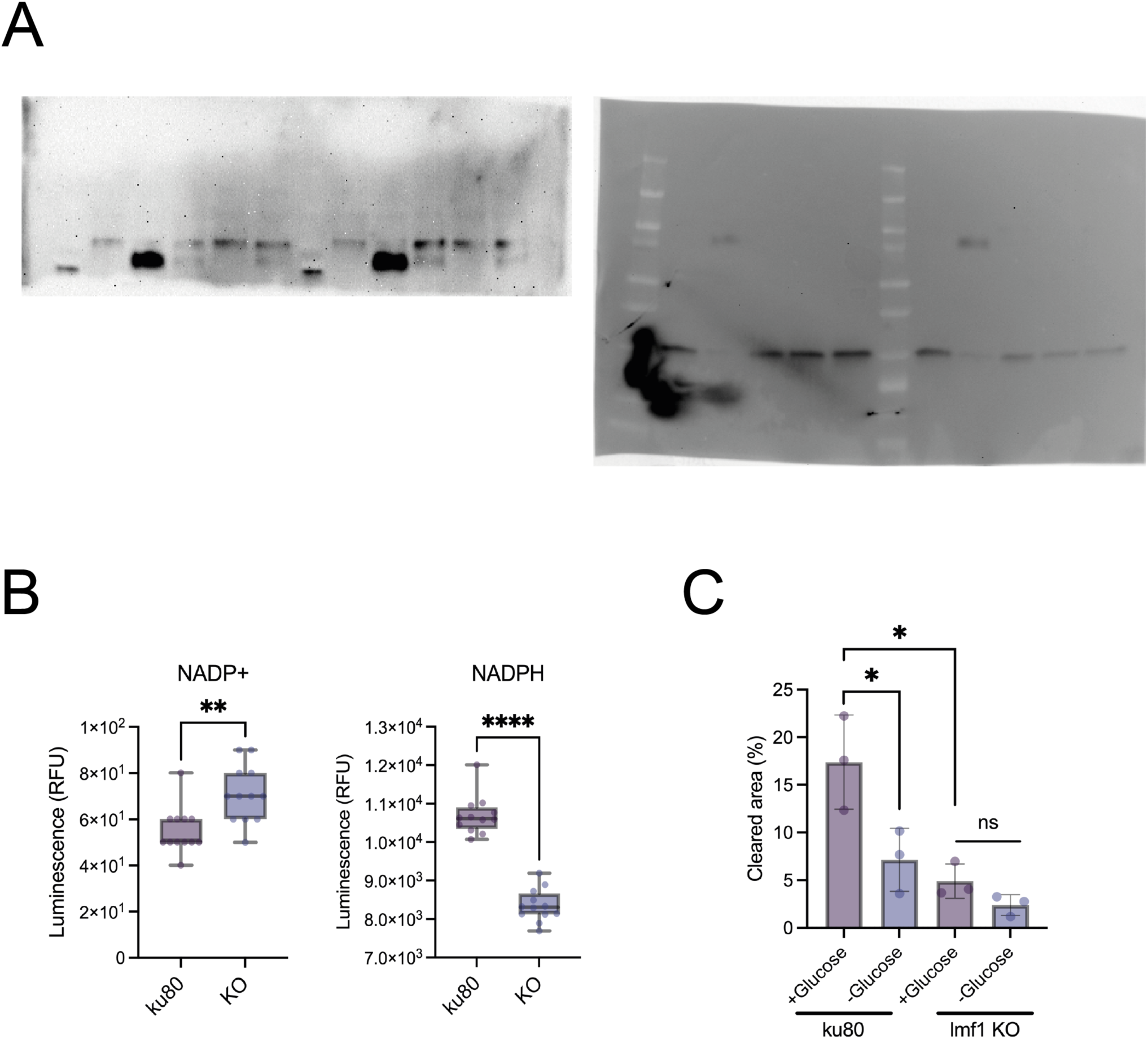
A) Uncropped original western blots. B) NADP+ and NADPH levels of Δku80 and Δlmf1 parasites. NADP(H) levels were measured using the NADP/NADPH-Glo Assay Kit (Promega) according to the manufacturer’s instructions. The experiment was performed in biological triplicate, with 4 technical replicates per replicate (n=12). Data are means ± s.d.. Statistical analysis is a Two-tailed unpaired t-test; **** P<0.0001, ** P<0.01; ns not significant (P>0.05). C) Quantification of the total cleared area per well of Δku80 and Δlmf1 cell lines grown in the presence (+) or absence (-) of glucose after a 5-day incubation period. Plaque assays were performed in three biological replicates. Data are means ± s.d.. Statistical analysis is a Two-tailed unpaired t-test; * P<0.05; ns = not significant.

## Supplemental Datasets

**Supplemental dataset 1.** RNA sequencing results for Δlmf1 vs. Δku80.

**Supplemental dataset 2.** RNA sequencing results for Δlmf1 vs. complemented (Δlmf1 + *LMF1-HA*).

**Supplemental dataset 3.** RNA sequencing results for complemented (Δlmf1 + *LMF1-HA*) vs Δku80.

